# How PTEN mutations degrade function at the membrane and life expectancy of carriers of mutations in the human brain

**DOI:** 10.1101/2023.01.26.525746

**Authors:** Hyunbum Jang, Jiaye Chen, Lilia M Iakoucheva, Ruth Nussinov

## Abstract

PTEN dysfunction, caused by loss of lipid phosphatase activity or deletion, promotes pathologies, cancer, benign tumors, and neurodevelopmental disorders (NDDs). Despite efforts, exactly how the mutations trigger distinct phenotypic outcomes, cancer or NDD, has been puzzling. It has also been unclear how to distinguish between mutations harbored by isoforms, are they cancer or NDDs-related. Here we address both. We demonstrate that PTEN mutations differentially allosterically bias P-loop dynamics and its connection to the catalytic site, affecting catalytic activity. NDD-related mutations are likely to sample conformations present in the wild-type, while sampled conformations sheltering cancer-related hotspots favor catalysis-prone conformations, suggesting that NDD mutations are weaker. Analysis of isoform expression data indicates that if the transcript has NDD-related mutations, alone or in combination with cancer hotspots, there is high prenatal expression. If no mutations within the measured days, low expression levels. Cancer mutations promote stronger signaling and cell proliferation; NDDs’ are weaker, influencing brain cell differentiation. Further, exon 5 is impacted by NDD or non-NDD mutations, while exon 7 is exclusively impacted by NDD mutations. Our comprehensive conformational and genomic analysis helps discover how same allele mutations can foster different clinical manifestations and uncovers correlations of splicing isoform expression to life expectancy.

## Introduction

Tumor suppressor phosphatase and tensin homologue (PTEN) acts as a dual-specific protein and lipid phosphatase, suppressing cell growth and survival (Tu et al., 2020). A major role of PTEN is the negative regulation of phosphoinositide 3-kinase (PI3K)/phosphoinositide-dependent protein kinase 1 (PDK1)/protein kinase B (AKT)/mammalian target of rapamycin (mTOR) signaling through dephosphorylation of the signaling lipid phosphatidylinositol 3,4,5-trisphosphate (PIP_3_) to phosphatidylinositol 4,5-bisphosphate (PIP_2_) (Georgescu, 2010). Dysfunction of PTEN due to somatic and germline genetic variations is associated with many different disease phonotypes. While somatic mutation of PTEN after conception is often associated with human cancers including glioblastomas and endometrial carcinomas (Koboldt et al., 2021; Sansal and Sellers, 2004), germline mutations (in egg or sperm cells) lead to neurodevelopmental disorders (NDDs) such as macrocephaly/autism syndrome (OMIM # 605309) (Busch et al., 2019; Morris-Rosendahl and Crocq, 2020) and PTEN hamartoma tumor syndrome (PHTS) (Abkevich et al., 1995). PHTS is a rare inherited syndrome characterized by a benign noncancerous tumor-like cell growth, including Cowden syndrome (CS) and Bannayan-Riley-Ruvalcaba syndrome (BRRS) (Cummings et al., 2022; Pilarski et al., 2013). Individuals with CS and BRRS open have macrocephaly, a non-tumoural phenotype. Further, individuals with PHTS genetic disorder have increased risk for certain types of cancer and autism spectrum disorder (ASD) (Butler et al., 2005; Buxbaum et al., 2007; Tan et al., 2012; Yehia et al., 2022).

The *PTEN* gene encodes the second most frequently mutated protein in human cancer followed by *TP53* (Yin and Shen, 2008). The most common PTEN mutations are nonsense, frameshift, and deletion/insertion (Bonneau and Longy, 2000). They likely result in premature termination of translation, which would decrease the level of PTEN protein in the cell. In addition, a considerable number of PTEN mutations are missense point substitutions (Serebriiskii et al., 2022) that may result in loss of protein function including reduced catalytic activity and protein stability at the membrane. Missense mutations including indel mutations are commonly located at the phosphatase domain, while nonsense mutations including truncation and frameshift are largely found in the C2 domain (Bonneau and Longy, 2000; Serebriiskii *et al*., 2022). In addition to the mutations, posttranslational modifications (PTMs) on the C-terminal tail through the phosphorylation of Ser/Thr cluster (Ser380, Thr382, Thr383, and Ser385) (Figure 1A) hamper PTEN’s cellular membrane localization, silencing its catalytic activity (Bolduc et al., 2013; Dempsey et al., 2021; Henager et al., 2016). In human malignancies, premature terminations, missense and nonsense mutations, frameshift mutations with frame deletion, PTMs including phosphorylation, ubiquitination, oxidation of active-site, and acetylation elevate uncontrolled PI3K-stimulated cell growth and survival (Alvarez-Garcia et al., 2019; Kotelevets et al., 2020; Meng et al., 2016; Singh and Chan, 2011; Song et al., 2012; Xia et al., 2020; Xu et al., 2014; Zhang et al., 2020).

**Figure 1.**
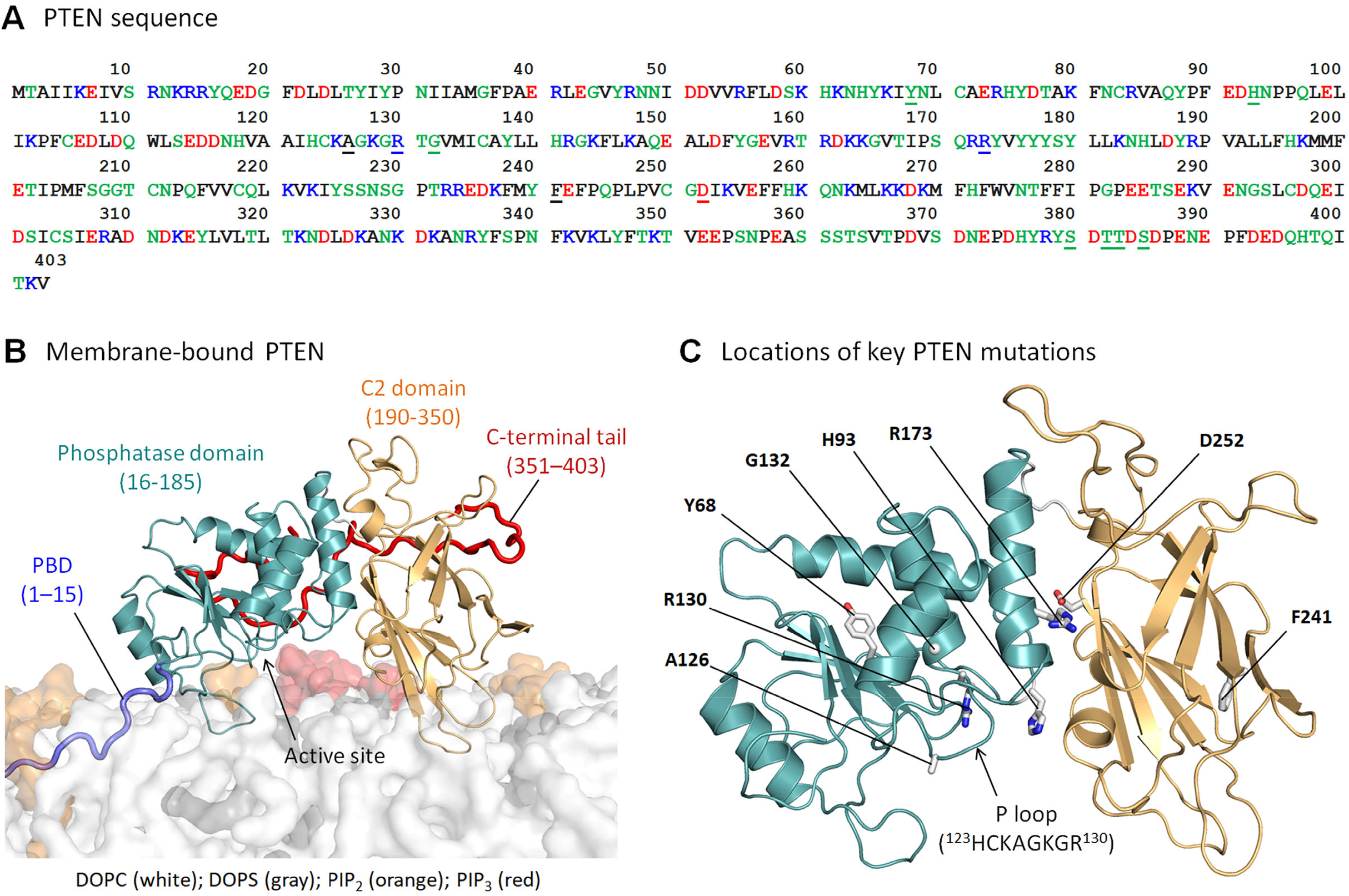
Sequence and mutations of PTEN. (A) The sequence of PTEN. In the sequence, the underlined residues highlight the mutation sites in the phosphatase and C2 domains, and the phosphorylated sites in the serine-threonine cluster of C-terminal tail. The residue letters are colored based on their amino acid types. (B) *In silico* model of the full-length PTEN interacting with the anionic lipid bilayer composed of DOPC:DOPS:PIP_2_:PIP_3_ (32:6:1:1, molar ratio). (C) Mapping of the residues for the mutations on the PTEN structure showing the phosphatase and C2 domains. P loop containing the catalytic signature motif ^123^HCxxGxxR^130^ is marked.

Although a number of experimental studies have demonstrated loss of PTEN lipid phosphatase activity due to mutations, mechanistic details of the mutations and the structural features of the mutant proteins at atomic resolution are still unknown. Here, comprehensive computational studies using molecular dynamics (MD) simulations were performed for PTEN mutants at an anionic lipid bilayer, composed of the phospholipids, phosphatidylcholine (PC) and phosphatidylserine (PS), and the phosphoinositides, PIP_2_ and PIP_3_ (Figure 1B). We only considered PTEN with the missense point substitutions, since proteins with the nonsense mutations and premature terminations are not amenable to MD simulations. Eight missense mutations of PTEN were considered: six in the phosphatase (Y68H, H93R, A126T, R130Q, G132D, and R173C) and two in the C2 (F241S and D252G) domains (Figure 1C). The types of mutations selected for the residues were with different chemical properties, ensuring that structural integrity of protein can be observable due to the mutations within the simulation time. Among them, Y68H is in the core of phosphatase domain and H93R is in the WPD loop (residues 88-98). The A126T, R130Q, and G132D mutations occur in and near the P loop (residues 123-130) with the catalytic signature motif, ^123^HCxxGxxR^130^ (where x is any amino acid). R173C is located at the interface between the phosphatase and C2 domains. For the C2 mutations, F241S is in the β-sandwich of the C2 domain and D252G is located at the interface between two major domains. Our studies indicate that the PTEN mutants can effectively absorb the anionic lipid bilayer, similar to wild-type PTEN. However, the mutations significantly reduce protein stability and hinder substrate recruitment. The dynamics of the P loop were restrained due to the strong allosteric signals from the mutation sites, which would affect the PTEN’s catalytic activity.

Our results underscore the merit of detailed structural and functional mechanisms of PTEN with mutations at the membrane, point how they may help resolve the enigma of how same-protein mutations can promote different pathologies, cancer versus NDDs, and a way to help determine their outcome. The sampled conformations of mutants harboring a mutation associated with an NDD resemble those of the wild-type protein. In contrast, conformations sampled by variants associated with cancer hotspots differ and indicate more potent catalytic activation. This supports the hypothesis that a key difference between cancer and NDDs mutations is mutation strength (Nussinov et al., 2022b; c). A strong activating mutation promotes cell proliferation, a weak/mild mutation promotes differentiation. This suggests that mutation strength, as manifested in the biased conformational sampling that the mutant favors can be harnessed as a feature in identifying mutations connected with the distinct clinical manifestation, cancer or NDD, assisting in early diagnosis.

NDDs emerge during embryonic brain cell development, suggesting that in addition to mutations, the level of prenatal gene expression plays a vital role. We analyzed prenatal and postnatal expression levels of isoforms harboring NDD (macrocephaly/ASD)-related mutations alone or in combination with cancer mutations. All mutant-harboring isoforms were highly expressed in the prenatal time window, dropping following birth; if no mutations within the measured life span, lower prenatal expression. Cancer development results from multiple (more than one hotspot) mutations, emerging sporadically during life. NDDs mutation carriers have higher chances of cancer emergence, suggesting that NDDs-related mutations can combine with cancer mutations. If they reside at adjacent chromosomal regions, deletions/insertions can also infringe both.

Our analysis helps learning how same allele mutations can abet different clinical manifestations and uncovers correlations of splicing isoform expression with life expectancy. It observes that splicing isoforms that do not carry exon 5 are exclusively impacted by the NDD mutations, F241S and D252G. On the other hand, variants carrying exons 5 and 7 can be highly correlated with increased lifetime risk for certain types of cancer. Individuals afflicted with NDDs are known to have increased risk of cancer, in schizophrenia as much as 50% probability (Nordentoft et al., 2021). It is also high in e.g., autism (Liu et al., 2022a), and in intellectual disability (Achterberg et al., 1978; Liu et al., 2021). Our work also offers guidelines for identification of cancer and NDD mutational variants. If the transcript harbors unknown mutation types, they can be differentiated by their strengths; cancer mutations tend to be stronger, with higher signaling levels; NDD’s weaker, with moderate signaling. To differentiate between the mutations, statistics and atomistic simulations can help, although applying MD on a large scale is demanding. Sampling could be accelerated. However, the challenge in accelerated conformational sampling is to have it sensitive to sequence alterations.

## Results

A full-length PTEN contains 403 amino acids (Lee et al., 1999), consisting of the N-terminal PIP_2_- binding domain (PBD, residues 1-15), the phosphatase domain (residues 16-185), the C2 domain (residues 190-350), and the carboxy-terminal tail (CTT, residues 351-403) (Figure 1). The CTT includes the PDZ binding motif (PDZ-BM, ^401^TKV^403^) at the C-terminal end. For catalysis, the phosphatase domain provides three critical catalytic residues in the active site; Asp92 in the WPD loop, and Cys124 and Arg130 in the P loop. We performed MD simulations on eight different PTEN mutants interacting with an anionic lipid bilayer composed of PC, PS, PIP_2_, and PIP_3_. The initial configuration of PTEN mutants at the membrane is the “open-open” conformation (Malaney et al., 2013; Rahdar et al., 2009; Ross and Gericke, 2009), reflecting the relaxed PTEN conformation at the anionic lipid bilayer as observed in the wild-type case (Jang et al., 2021; Nanda et al., 2015; Shenoy et al., 2012). All PTEN mutants stably anchored in the anionic lipid bilayer. As observed in the wild-type PTEN system with the same lipid compositions (Jang *et al*., 2021), the probability distribution functions of membrane contacts of the protein residues point to five loops that are responsible for the membrane association (Figure 1– figure supplement 1). The peaks in the distribution indicate PBD-pβ1(^19^DGFDL^23^) and pβ2-pα1 (^41^RLEGVYR^47^) loops in the phosphatase domain, and cβ1-cβ2 (^205^MFSGGTC^211^), CBR3 (^260^KQNKMLKKDK^269^), and Cα2 (^327^KANKDKANR^335^) loops in the C2 domain. In addition to the PBD, the two positively charged loops, pβ2-pα1 and CBR3 loops, one from the phosphatase domain and the other from the C2 domain, are major membrane-binding interfaces of PTEN mutants. As observed in the wild-type systems, similar profiles of the distributions of the helix tilt angles for the helices in the phosphatase domain of PTEN mutants (Figure 1–figure supplement 2) suggest that membrane absorption and orientation of the protein are highly affected by the lipid compositions in the bilayer.

### Y68H in the core of phosphatase domain

In wild-type PTEN, Tyr68 in pβ3 forms an aromatic cluster with Tyr88 in pβ4 and Phe104 in pα3. In the Y68H mutant, the point substitution disrupts this cluster (Figure 2A), resulting in the destruction of the salt bridge between Lys66 in pβ3 and Asp107 in pα3 (Figure 2B). The membrane absorption of the pβ2-pα1 loop in the phosphatase domain seems to be weaker than that of the other mutants and wild-type system (Figure 1–figure supplement 1). The disruptions of key residue interactions cause a conformational change in the phosphatase domain, yielding a loosely packed core structure. This provides room for the mutant residue His68 to rotate its aromatic ring. The periodic fluctuations in the distance between HD1 at the ring and HB2 at C_β_ atom indicate the rotation of His68 aromatic sidechain (Figure 2C). In comparison with wild-type PTEN, no rotation of the aromatic ring of Tyr68 is monitored. To observe how the mutation allosterically affects the conformation of the active site, we identified the signal propagation pathways through the protein by calculating the dynamic correlated motion among residues using the weighted implementation of suboptimal paths (WISP) algorithm (Van Wart et al., 2014). A number of optimal and suboptimal pathways were generated between the source residue, His68 (or Tyr68 for wild type), and the sink resides, Cys124 and Arg130, in the P loop (Figure 2D). The allosteric signal propagations through the protein illustrate that the mutant residue His68 is dynamically correlated with the P loop residues, Cys124, Lys125, Arg130, and Thr131. The strong allosteric signals due to the mutation transmitting through the active site constrain the P loop to move upwards from the bilayer surface (Figure 2E). In marked contrast to the mutant system, the allosteric signal nodes of Lys125 and Thr131 are absent in the signal propagation pathways from the wild-type residue Tyr68, implicating weak allosteric coupling to the P loop. For Y68H, the allosteric restraint on the P loop with the shifted conformation hampers the catalytic residue Arg130 recruitment of the substrate PIP_3_ (Figure 2F), which can lead to reduced catalytic activity.

**Figure 2.**
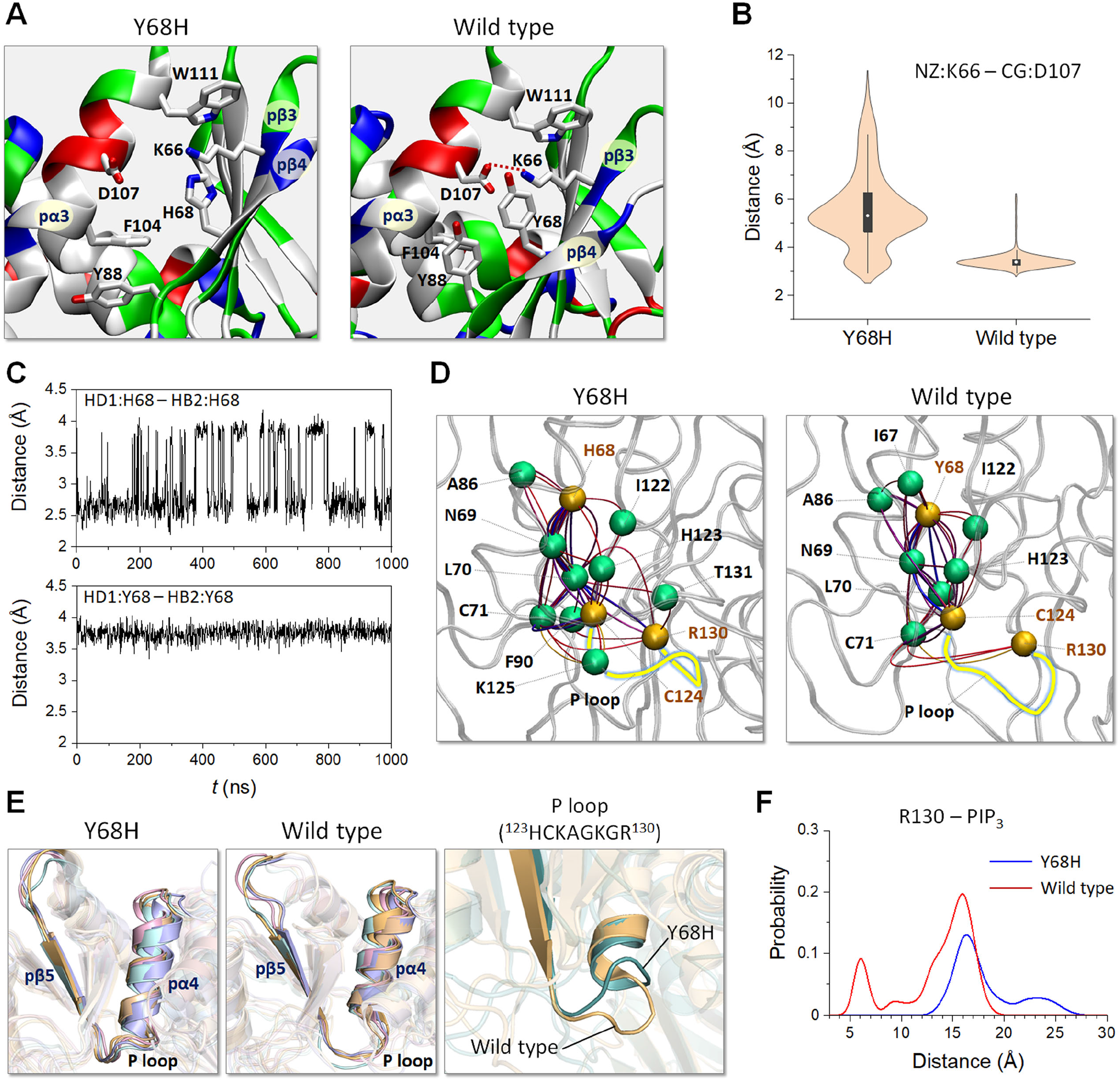
Y68H in the core of phosphatase domain. (A) The best representative conformation from the ensemble clusters highlighting the mutation site of Y68H. The wild-type PTEN is shown for comparison. In the cartoons, residues are colored based on their amino acid types. In wild-type PTEN, red dotted line denotes a salt bridge. (B) Violin plots representing the atomic pair distance between NZ of Lys66 in pβ3 and CG of Asp107 in pα3 for Y68H and wild-type PTEN. (C) The time series of atomic pair distances between HD1 and HB2 of His68 for Y68H (upper panel) and Tyr68 for wild-type PTEN (lower panel). (D) The allosteric pathways between the mutation site and P loop. The source residues are His68 for Y68H and Tyr68 for wild-type PTEN, and the sink residues are Cys124 and Arg130 for both proteins. Yellow beads represent the source and sink residues, and green beads denote the allosteric signal nodes. The blue lines represent the shortest allosteric paths. The P loop is colored yellow. (E) Superimpositions of the top five representative conformations of P loop for Y68H (left panel) and wild-type PTEN (middle panel). Superimposition of the first representative conformations of P loop from Y68H and wild-type PTEN (right panel). (F) The probability distribution of the center of mass distances between the guanidine group of Arg130 and the phosphate group in the inositol of PIP_3_ for Y68H and wild-type PTEN.

### H93R in the WPD loop

The P loop is highly basic, containing three basic residues that facilitate the recruitment of the acidic substate PIP_3_ to the catalytic pocket. For catalysis, three catalytically significant residues, Cys124 and Arg130 in the P loop and Asp92 in the WPD loop, align to coordinate PIP_3_ at the active site. The WPD loop in a closed conformation can bring Asp92 in the coordination, leading to high catalytic activity (Brandao et al., 2012). A point substitution H93R in the WPD loop amplifies the positively charged nature of the active site (Figure 3). The location of the WPD loop with respect to the P loop is comparable to the wild type (Figure 3–figure supplement 3), suggesting that H93R preserves the closed loop conformation. However, the mutant residue Arg93 increases the interaction with the substate PIP_3_, which seems to block the migration of the substrate to the catalytic site residues. This additional membrane interaction might be correlated with the absence of the membrane contact of the Cα2 loop in the C2 domain (Figure 1–figure supplement 1A).

**Figure 3.**
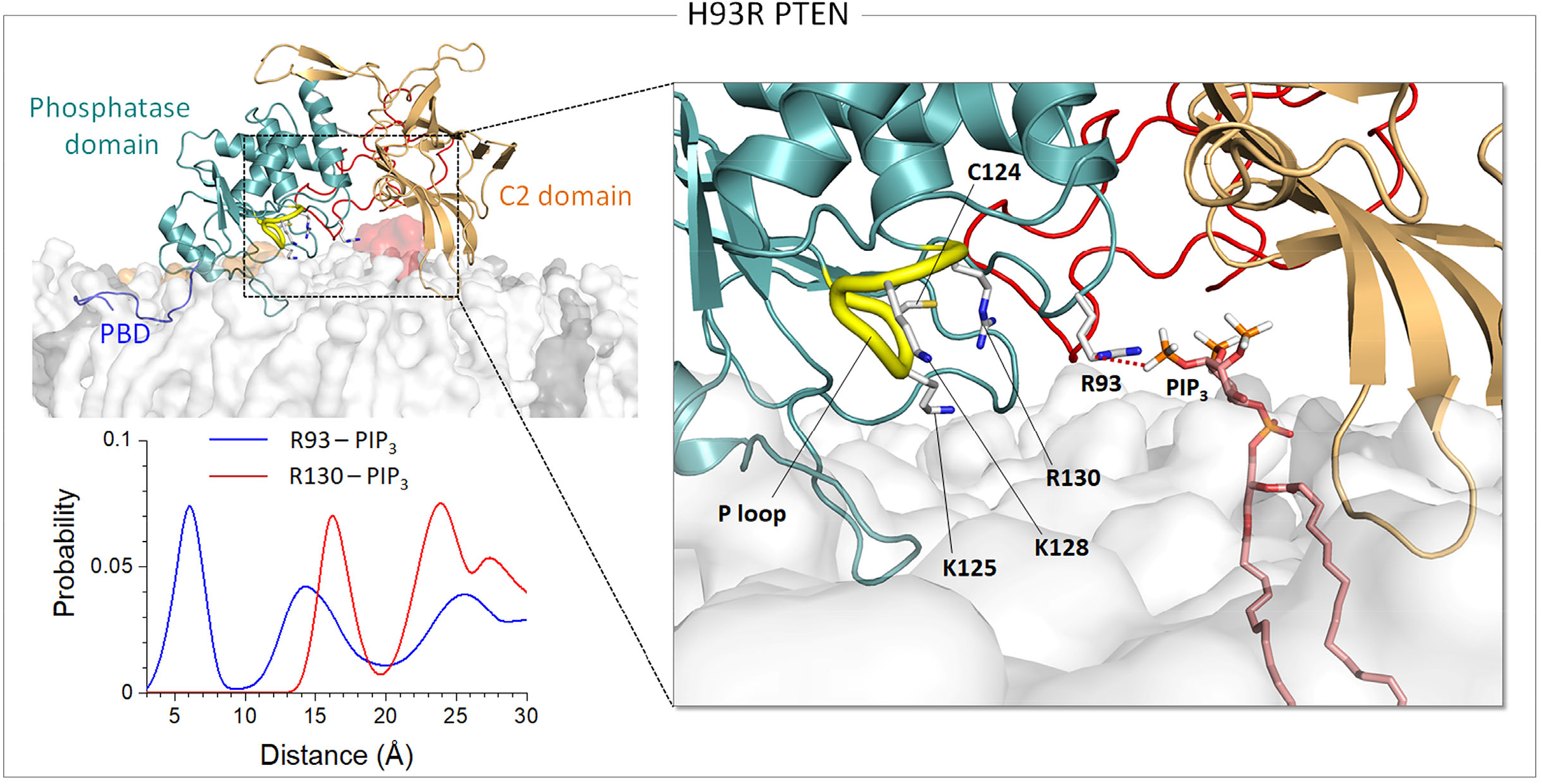
H93R in the WPD loop. Snapshot representing the best representative conformation from the ensemble clusters for H93R in the anionic lipid bilayer (top left). Highlight showing the interaction of the mutant residue Arg93 with PIP_3_ (right). The probability distribution of the center of mass distances between the guanidine groups of Arg93, or Arg130 for comparison, and the phosphate group in the inositol of PIP_3_ for H93R (bottom left).

### A126T, R130Q, and G132D in the P loop

The P loop contains the catalytic signature motif ^123^HCKAGKGR^130^, suggesting that any mutation of P loop residue can alter the loop conformation. Our data illustrate that the direct P loop mutations, A126T and R130Q, and G132D nearby the P loop, induce a collapsed loop conformation (Figure 4A). In contrast, it was found that an extended (or relaxed) conformation of the P loop is populated for wild-type PTEN when the anionic bilayers contain both PIP_2_ and PIP_3_ (Jang *et al*., 2021). Although our mutant systems contain the same phosphoinositide lipids, they yield the collapsed P loop conformation regardless of the lipid composition. Interestingly, both A126T and R130Q mutants show an open conformation of the WPD loop with increased distance from the P loop as compared to the wild type (Figure 4B). However, the G132D mutant maintains a closed conformation of the WPD loop with the distance from the P loop comparable to wild-type PTEN, suggesting that G132D exhibits weaker mutational effect compared to the other mutations.

**Figure 4.**
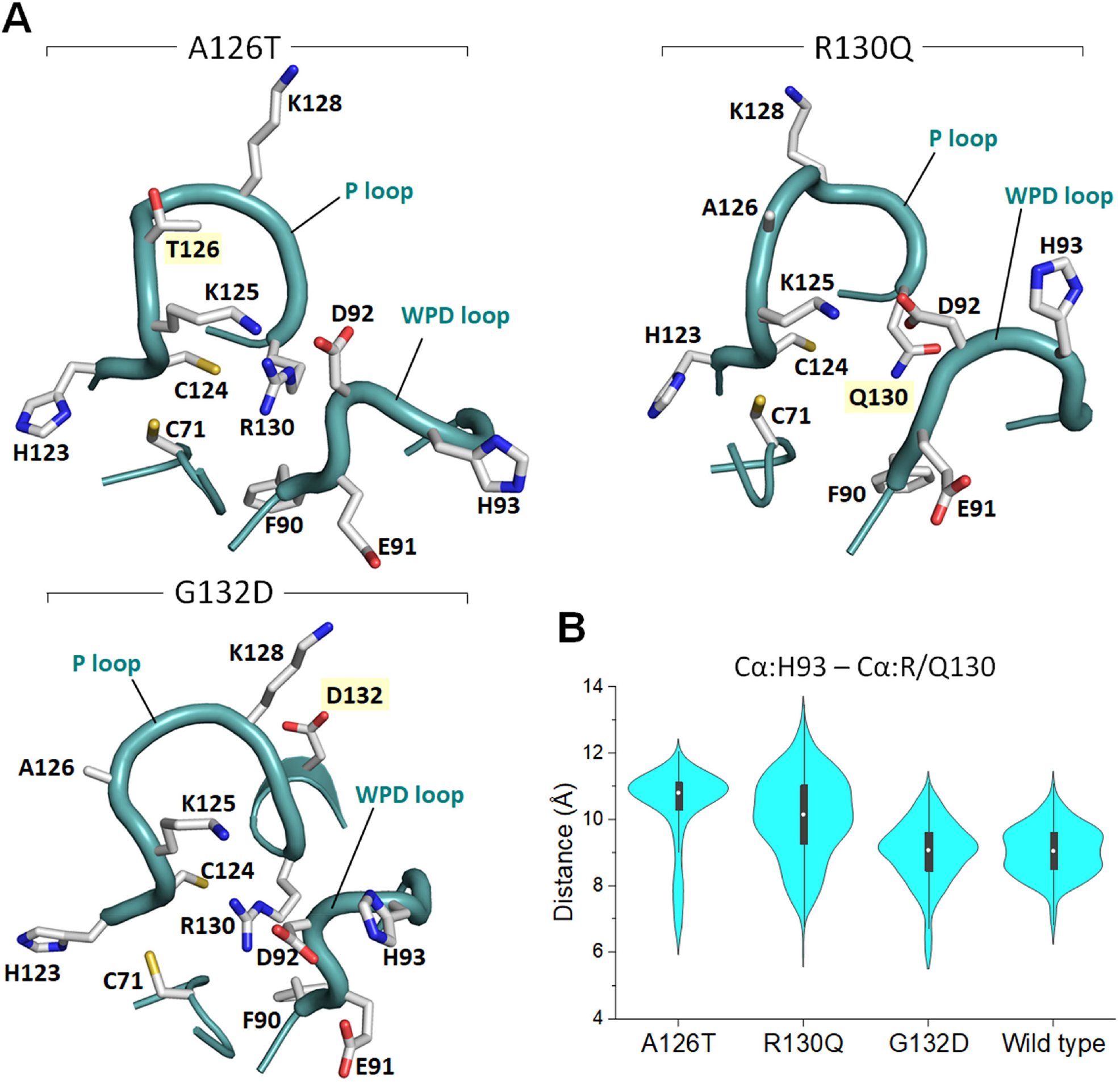
A126T, R130Q, and G132D in the P loop. (A) The conformations of P loop and WPD loop for A126T, R130Q, and G132D. Key residues are marked, and the mutated residues are marked with yellow background. (B) Violin plots representing the atomic pair distance between Cα of His93 in the WPD loop and Cα of Arg130 (Gln130 for R130Q) in the P loop for A126T, G132D, and wild-type PTEN.

For catalysis, PTEN requires residual water molecules around the sidechains of Cys124 and Arg130 at the active site in the process of hydrolysis to release the phosphate group from Cys124 after transferring it from PIP_3_ (Brandao *et al*., 2012). To delineate the catalytic activity in the mutant systems, we calculated the three-dimensional water density map in the region of the phosphatase domain (Figure 5). Compared to the wild-type system, low probability of water around the catalytic residues indicates that the active sites of A126T, R130Q, and G132D mutants are largely dehydrated. The severe dehydration in the active site of R130Q suggests that the mutational effect may be stronger than the other mutants. For R130Q, changes in the helix tilt angles for pα3 and pα5 are apparent when compared to wild-type PTEN (Figure 1–figure supplement 2A).

**Figure 5.**
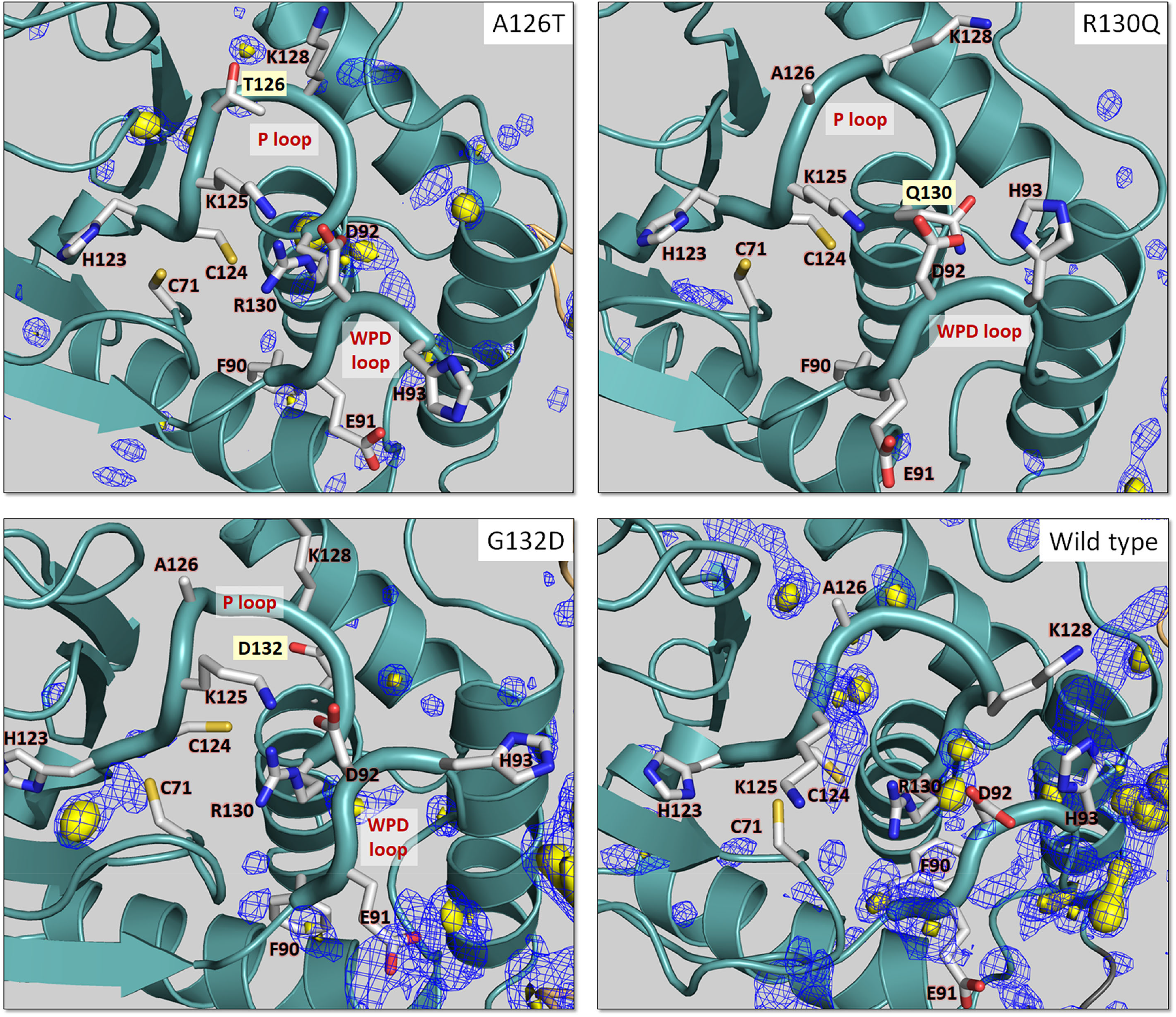
Water density in the active site. Three-dimensional water density map with probabilities *P* = 0.5 (yellow surface) and *P* = 0.4 (blue mesh) for A126T, R130Q, and G132D. Also showing wild-type PTEN for comparison. The protein structures depict the best representative conformation from the ensemble clusters. The mutated residues are marked with yellow background.

### R173C at the interface

In wild-type PTEN, Arg173 in pα6 of the phosphatase domain is important for maintaining the interdomain interaction at the interface between the phosphatase and C2 domains. It forms a strong salt bridge with Asp324 in the cβ7-cα2 loop, which induces the interdomain π-π stacking between Tyr177 in pα6 and Phe279 in cα1 (Figure 6A). In the R173C mutant, the absence of the salt bridge actuates the destabilization of the interface, resulting in the disruption of the π-π stacking. The removal of these key residue interactions increases the interdomain distance at the mutation site (Figure 6B). However, the opposite site of the interface is still maintained by the hydrophobic interaction between Pro95 in the WPD loop and Trp274 in cβ6, and an additional salt bridge formation between Gln97 in the WPD loop and Asp252 in cβ5. This unbalanced interaction in the interface induces the rotation of the C2 domain with respect to the phosphatase domain (Figure 6C), causing the loss of the membrane contact of the Cα2 loop in the C2 domain (Figure 1–figure supplement 1A). The allosteric signaling pathways from the mutant residue Cys173 to the catalytic residue Arg130 seem to be stronger than those from the wild-type residue Arg173 (Figure 6D). This suggests that the R173C mutant allosterically constrains the P loop through the multiple shortest optimal pathways. The allosteric restraint on the P loop changes the loop conformation that moves upwards from the bilayer surface as observed in Y68H (Figure 6E). The shifted P loop that the location is highly elevated from the bilayer surface and adopts a collapsed loop conformation (Figure 6F), which induces the WPD loop open conformation. We observed that the substrate PIP_3_ is populated in the region of the C2 domain. The failure of the R173C mutant to recruit PIP_3_ by Arg130 (Figure 6G) indicates that it has a reduced catalytic activity.

**Figure 6.**
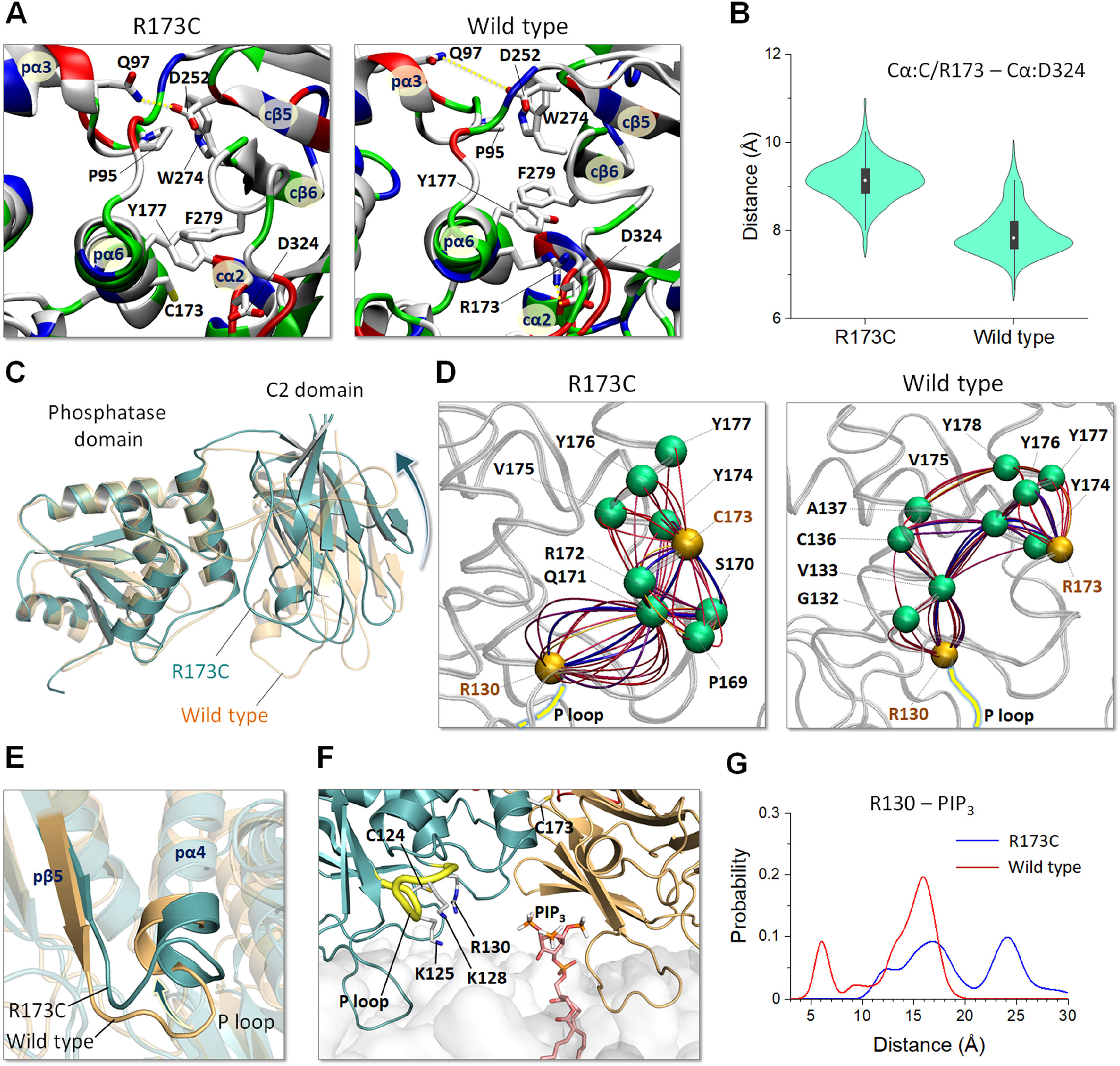
R173C at the interface. (A) The best representative conformation from the ensemble clusters highlighting the mutation site of R173C. Also showing wild-type PTEN for comparison. In the cartoons, residues are colored based on their amino acid types. Yellow dotted lines denote salt bridges. (B) Violin plots representing the atomic pair distance between Cα of Cys173 (Arg173 for wild-type PTEN) in pα6 and Cα of Asp324 in the cβ7-cα2 loop for R173C. (C) Superimposition of the first representative conformations of R173C and wild-type PTEN with respect to the phosphatase domain. (D) The allosteric pathways between the mutation site and P loop. The source residues are Cys173 for R173C and Arg173 for wild-type PTEN, and the sink residue is Arg130 for both proteins. Yellow beads represent the source and sink residues, and green beads denote the allosteric signal nodes. The blue lines represent the shortest allosteric paths. The P loop is colored yellow. (E) Superimposition of the first representative conformations of P loop from R173C and wild-type PTEN. (F) Snapshot representing the best representative conformation from the ensemble clusters for R173C. Highlight showing the interaction of PIP_3_ with the C2 domain. (G) The probability distribution of the center of mass distances between the guanidine group of Arg130 and the phosphate group in the inositol of PIP_3_ for R173C and wild-type PTEN.

### F241S and D252G in the C2 domain

F241S in cβ4 resides in the pocket of the β-sandwich of the C2 domain, forming a hydrophobic cluster. D252G in cβ5 occurs at the interface between the phosphatase and C2 domains, similar to R173C. As expected, both C2 mutations increase the fluctuations in the C2 domain as compared to wild-type PTEN (Figure 7A). However, averaged deviations of the key basic residues from the bilayer surface are markedly different between these two C2 mutants (Figure 7B). The profile of averaged deviations of F241S resembles that of wild-type PTEN, but that of D252G is distinct. F241S shows a relatively weak membrane absorption of the pβ2-pα1 loop (Figure 1–figure supplement 1B), and D252G alters the helix tilt angles for the helices in the phosphatase domain (Figure 1–figure supplement 2B). F241S destabilizes the hydrophobic core of the β-sandwich (Figure 7C), affecting the dynamic correlations of motions of the residues in the C2 domain. The allosteric signal propagations from the mutant residue Ser241 to the active site avoid the signal nodes in the hydrophobic core, while the allosteric signals from the wild-type residue Phe241 transmit through the signal nodes in the hydrophobic core of the β-sandwich (Figure 7D). F241S obtains a single optimal pathway that passes more allosteric signal nodes than the wild type, indicating less effective allosteric connection to the active site. In contrast, D252G exhibits strong allosteric connection to the active site (Figure 7E). The allosteric signals from the wild-type residue Asp252 propagate through the signal nodes at the interface, Pro95 and Trp274, and in the WPD loop, Glu91, Asp92, His93, Asn94, and Pro96. However, the allosteric signals transmitting through the WPD loop is missing in the D252G mutant. The loss of the hydrophobic interaction due to the mutation destabilizes the interface (Figure 7F) and increases the interfacial distance (Figure 7G). D252G shows the similar behavior as observed in R173C since both mutations occur in the same interface but in different side.

**Figure 7.**
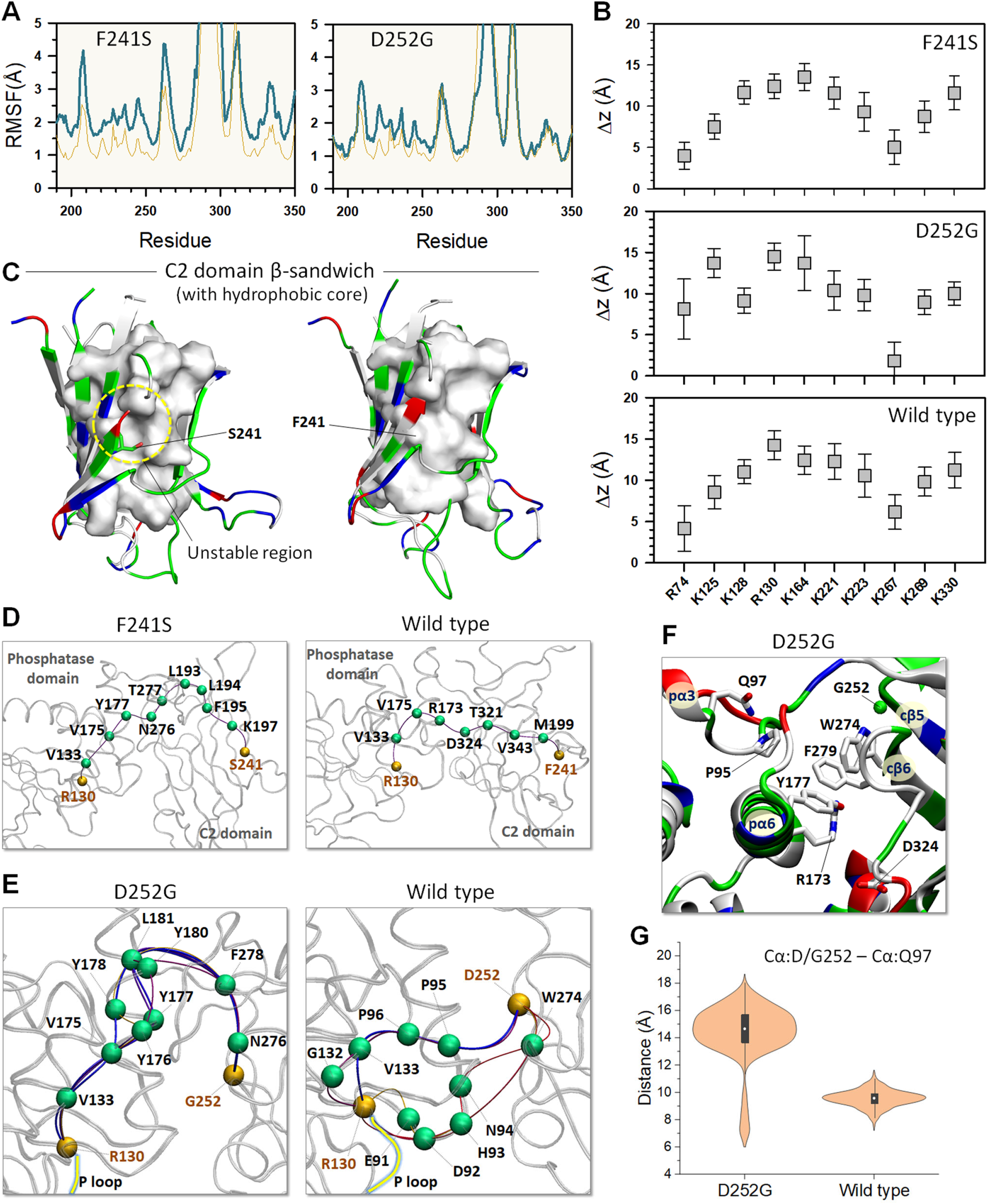
F241S and D252G in the C2 domain. (A) The root-mean-squared-fluctuations (RMSFs) of the C2 residues for F241S (left panel) and D252G (right panel). Thin orange lines represent the RMSF of wild type PTEN for comparison. (B) Averaged deviations of the amide nitrogen in the sidechains of Arg and Lys residues from the bilayer surface for the PIP_3_-favored residues in the phosphatase and C2 domains for F241S and D252G. Also showing wild-type PTEN for comparison. Error bars denote standard deviation. (C) Snapshot highlighting the hydrophobic core (surface representation in white) in the β-sandwich of C2 domain for F241S and wild-type PTEN. The protein structures depict the best representative conformation from the ensemble clusters. The allosteric pathways between the mutation site and P loop for (D) F241S and (E) D252G. In (D), the source residues are Ser241 and Phe241 for F241S and wild-type PTEN, respectively, and in (E) they are Gly252 and Asp252 for D252G and wild-type PTEN, respectively. The sink residue is Arg130 for all proteins. Yellow beads represent the source and sink residues, and green beads denote the allosteric signal nodes. The blue lines represent the shortest allosteric paths. The P loop is colored yellow. (F) The best representative conformation from the ensemble clusters highlighting the mutation site of D252G. In the cartoons, residues are colored based on their amino acid types. (G) Violin plots representing the atomic pair distance between Cα of Gly252 (Asp252 for wild-type PTEN) in cβ5 and Cα of Gln97 in the WPD for D252G.

### PTEN variants expressions for NDD vs. non-NDD

PTEN mutations are associated with various diseases including PHTS, cancer, and NDDs. Some PTEN mutations are exclusively expressed in a certain disease type, but mutations can share across both disease phenotypes. Here, the NDD-related mutations are H93R, F241S, and D252G that are exclusively responsible for macrocephaly/autism syndrome (Butler *et al*., 2005). The *PTEN* gene is located on the chromosome 10 and contains nine exons. The longest human PTEN splicing isoform is encoded by the transcript ENST00000371953, with exon 3 (Y68H) and exon 6 (R173C) being impacted by the non-NDD mutations, exon 7 (F241S and D252G) by the NDD mutations, and exon 5 (H93R, A126T, R130Q, and G132D) by both, the NDD and non-NDD mutations (Figure 8A).

**Figure 8.**
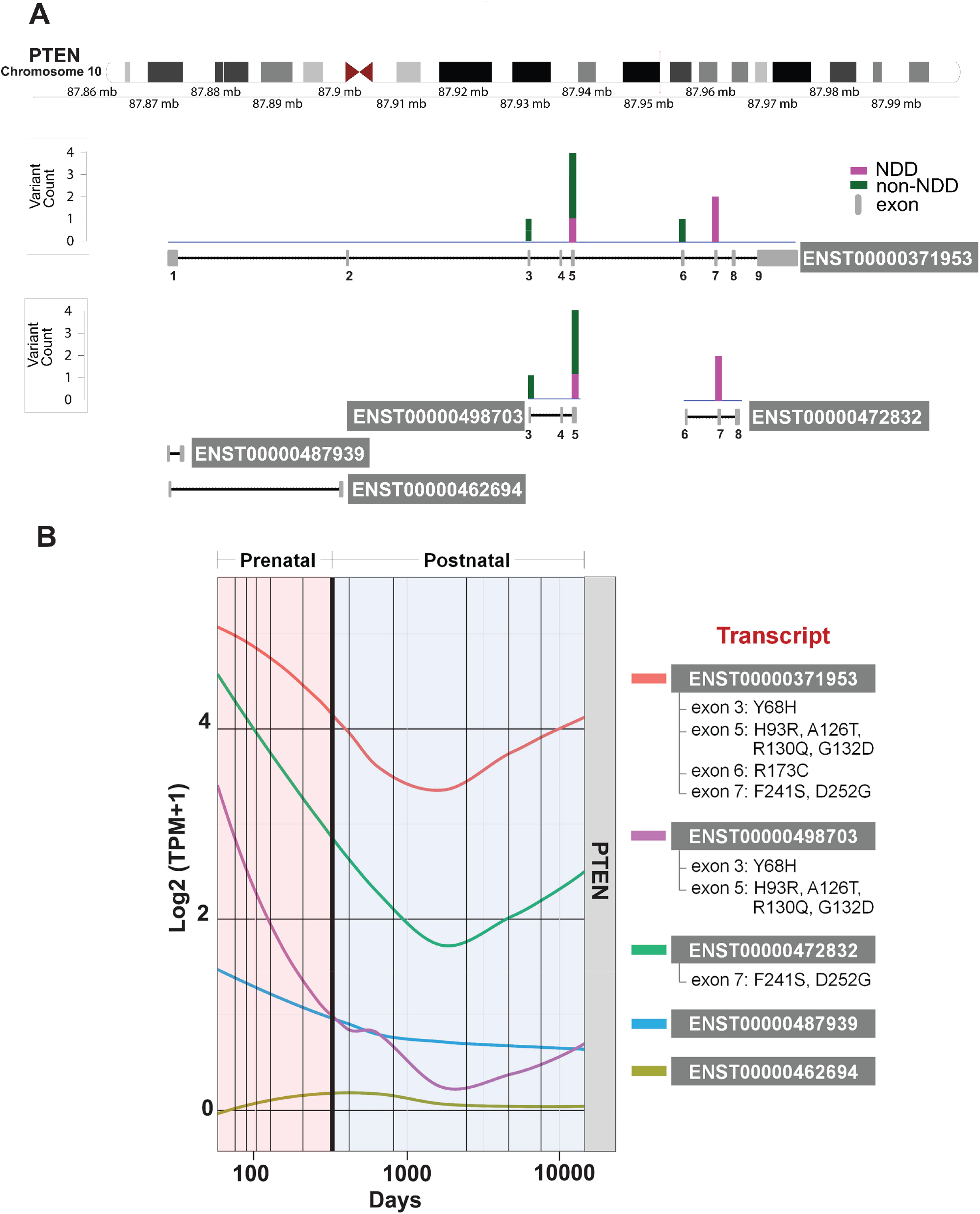
PTEN variants mapping and expression. (A) Mapping of variants implicated in neurodevelopmental disorders (NDDs, pink) and those from other diseases (green) to PTEN splicing isoforms. (B) Expression of PTEN isoforms in the developing human brain for which expression levels are available. The isoform expression data was quantified by the PsychEncode Consortium. Three PTEN isoforms (red, green and purple) are highly expressed prenatally, and their expression levels decrease after birth. PTEN isoforms and their associated exons and mutations are marked.

Two other transcripts (ENST00000498703 and ENST00000472832) are shorter isoforms that carry above combinations of mutations, except that exon 6 in the ENST00000472832 isoform (87952199-87952259, GRCh38.p13) is slightly (∼80 bp) shorter than the same exon in the ENST00000371953 isoform (87952118-87952259, GRCh38.p13) due to alternative splicing, and it therefore carries only two NDD mutations in its exon 7 (F241S and D252G) and is missing a non-NDD mutation R173C from exon 6. There are additional isoforms comprising exons 1 and 2 that do not carry any known disease risk mutations. We quantified expression levels of these five isoforms from the RNA-seq dataset of the developing human brain BrainSpan (Kang et al., 2011; Li et al., 2018), as we have previously described (Chau et al., 2021), and observed that three PTEN isoforms (ENST00000371953, ENST00000472832, and ENST00000498703) are highly expressed prenatally, and their expression levels decrease after birth (Figure 8B). The remaining two isoforms (ENST00000487939 and ENST00000462694) that are not impacted by mutations, are lowly expressed in the developing brain. Interestingly, the isoform ENST00000472832 with the shorter exon 6 that is exclusively impacted only by the NDD mutations, is the second highly expressed PTEN isoform, which may have further implications for NDD biology.

## Discussion

Here, we considered six PTEN mutations in the phosphatase domain (Y68H, H93R, A126T, R130Q, G132D, and R173C) and two in the C2 domain (F241S and D252G). Our studies demonstrate that the PTEN mutants retain the wild-type capability of the membrane absorption to the anionic lipid bilayer (Han et al., 2000). However, the dynamics of the P loop, the WPD loop conformation, the hydration of the active site, and the substrate recruitment were greatly affected by the mutations. Y68H is associated with CS, BRRS, and glioblastoma, which is known to be affected by the loss of phosphatase activity and protein stability (Han *et al*., 2000; He et al., 2011; Marsh et al., 2001; Post et al., 2020; Tsou et al., 1998). In our simulations, Y68H disrupted the core of the phosphatase domain and allosterically constrained the P loop, which hinders the recruitment of the substrate PIP_3_. The NDD-related mutation H93R is responsible for macrocephaly/autism syndrome, displaying a modest loss of catalytic activity (Fricano-Kugler et al., 2018; Redfern et al., 2010; Rodriguez-Escudero et al., 2011). In our structural model, H93R in the WPD loop hijacked the substate PIP_3_, interrupting the catalytic site residues recruitment of the substate for catalysis. But the mutant protein preserved the closed WPD loop conformation. For the P loop mutations, A126T is found in endometrial and ovarian carcinomas (Valtcheva et al., 2017), and R130Q is shared by CS and endometrial carcinoma (Han *et al*., 2000; Serebriiskii *et al*., 2022). In our simulations, these mutations yielded a collapsed conformation of the P loop, resulting in the loss of contact with the WPD loop. G132D near the P loop, which is associated with endometrial carcinoma and ASD (Chao et al., 2020; Post *et al*., 2020), also exhibited the collapsed P loop conformation but preserved the closed WPD loop conformation. We observed that the PTEN mutations in the P loop, or nearby, cause dehydration in the active site, where water molecules are important for hydrolysis to release the phosphate group from the active site (Brandao *et al*., 2012). R130Q exhibited more severe dehydration than the other mutants. At the interface between the phosphatase and the C2 domains, R173C is associated with cancer, such as glioblastoma and endometrial carcinoma (Han *et al*., 2000; Shan et al., 2020). We found that R173C disrupts the domain-domain interaction, allosterically biasing the P loop dynamics. Similar behavior was observed for the C2 mutation D252G at the interface. However, the other C2 mutation, F241S in the β-sandwich of C2 domain, exhibited less effective allosteric connection to the catalytic site than that observed in wild-type PTEN. Both NDD-related C2 mutations F241S and D252G are responsible for macrocephaly/autism syndrome (Fricano-Kugler *et al*., 2018; Mingo et al., 2018; Post *et al*., 2020; Rodriguez-Escudero *et al*., 2011; Spinelli et al., 2015).

Total loss of protein function can occur when PTEN has: (i) reduced protein expression due to truncation and (ii) PTM, i.e., C-terminal tail phosphorylation in solution. In these cases, PTEN is totally removed from the cell membrane, dismissing its catalytic activity (Bolduc *et al*., 2013; Dempsey *et al*., 2021; Henager *et al*., 2016). On the other hand, PTEN with missense mutations can effectively absorb the cell membrane, exhibiting function with reduced activity (Han *et al*., 2000). We characterized the structural integrity of how PTEN degrades its function at the membrane due to missense mutations. Our membrane bound PTEN mutants exhibited key structural features: (i) the phosphatase domain with reduced stability, (ii) the allosteric constraint on the P loop, (iii) the collapsed P loop, (iv) the dehydration of active site, and (v) the open conformation of WPD loop. Although the simulations cannot directly assay PTEN lipid phosphatase activity, the failure in the coordination of the substate PIP_3_ at the catalytic residues is a corollary of all the above structural features that lead to silencing PTEN catalytic activity. The phosphatase mutations, Y68H, A126T, R130Q, and R173C have all the above structural features induced by the mutations, suggesting that these proteins appear to exhibit a strong mutational effect. In contrast, the NDD-related H93R and F241S exhibit a weak mutational effect with few structural features by the mutations. Both cancer- and NDD-related G132D and only NDD-related D252G exhibit an intermediate mutational effect with the structural features by the mutations.

In our studies, the phosphatase mutations are associated with cancer, PHTS, and NDDs, while the C2 mutations are exclusively related to NDDs. Principal component analysis (PCA) of the sampled conformations found that the macrocephaly and ASD related mutations, H93R and F241S, favor sampling conformations present in wild-type PTEN (Figure 8–figure supplement 4). In contrast, the sampled conformations for the cancer and PHTS-related mutations, Y68H, A126T, and G132D, differ from those in wild-type PTEN. The interface mutations R173C and D252G favor sampling similar conformations. Interestingly, although the sampled conformations for R130Q can overlap those of the wild-type PTEN, the function of the mutant protein largely differs. We suspect that a key structural effect of the PTEN missense mutation at the membrane is an impact on the dynamics and conformation of the P loop. The strong PTEN mutations, Y68H and R173C, which are distant from the active site, constrain the P loop through a strong allosteric signal, while R130Q, the mutation directly on the P loop, strongly controls the loop conformation. It was reported that cancer or PHTS-associated mutations targeting the P-loop of PTEN resulted in complete loss of protein function (Rodriguez-Escudero *et al*., 2011).

These distinct structural features in PTEN mutations appear to correlate with mutation strength and timing of the expression of the transcripts that determine the cancer and NDD outcomes. PTEN contains nine exons, and its mutations largely occur in exon 5, followed by exon 7, 3, and 6 (Tan et al., 2011). Most missense mutations occur within the phosphatase domain, while the C2 domain mainly accommodates nonsense mutations. The largest exon 5 encodes the PTEN residues 84-164 including the catalytic signature motif, ^123^HCxxGxxR^130^. It was found that up to 40% of all germline mutations are located in exon 5 (Waite and Eng, 2002). The developing brain isoform expression data indicate that exon 5 is impacted by NDD or non-NDD mutations. Interestingly, we observed that PTEN splicing isoforms that do not carry exon 5 are exclusively impacted by the NDD mutations, F241S and D252G. We expect that the increased life expectancy of PTEN variants carrying exons 5 and 7 can be highly correlated to increased lifetime risk for certain types of cancer.

## Conclusions

PTEN, like other proteins in the signaling networks of the Ras superfamily and their associated regulatory proteins harbor mutations connected with cancer and with NDDs. As a phosphatase, PTEN is undruggable. Its associated interactome can be. Early diagnosis could help in ASD pharmacology. Identifying the mutations acting in cancer, NDD, or both has been challenging. The timing of the expression is a major determinant, during embryonic development or sporadic, throughout life in cancer. Here our data suggest that mutation strength is another crucial factor. To determine the mutation strength here we exploit the conformations sampled by the mutants. If the conformations are biased toward the wild type, we interpret the mutation as low/mild, acting in NDD. If they differ, adopting catalytically favored states, we label them as tending to strong hotspots. Strong mutations result in a larger population of active molecules, thus stronger signals reaching the cell cycle to promote proliferation (Nussinov et al., 2022a; d). We suggest sampling as a general approach toward defining the likelihood of mutations to act in distinct pathology in diagnosis. The atomistic MD simulations used here are limited by molecular size, and the number of proteins and mutations. Accelerated MD can be applied on a broader scale. It could also be employed as a first step in sequence sensitive, deep modeling (Strokach and Kim, 2022). We expect that other proteins bearing NDD connected mutations also display biased conformations.

MD simulations are a powerful tool to gain insight into the molecular behavior of proteins, wild type, and mutants. However, in the living cell, the conformational behavior is not stand-alone, and the mutant behavior is insufficient in determining cell transformation (Nussinov *et al*., 2022a; d). In addition to the mutation strength, determinants of signal strength include mechanisms that can block or enhance the signal, the types, and locations of additional mutations, and critically, the expression levels of the respective isoforms, and of cross-talking proteins in the pathway that regulate the protein variants. Signal levels vary across cell types, states, and time windows, with chromatin structure and alternative splicing playing key roles. A strong activating mutation can be constrained by low expression level, and a weak/moderate mutation can be strengthened by high expression. Considering the spatio-temporal isoform expression in relevant tissues and cell types in conjunction with mutations can help unravel the molecular mechanisms driving human disease. Here, the expression levels of splicing isoforms harboring NDD, and mixed NDD/cancer mutations are elevated at the prenatal stage, dropping following birth. The mapping of these mutations on the respective exons and the presence of the exons in the isoforms, can be among the factors foretelling life expectancy.

## Materials and methods

### Construction of full-length PTEN protein with mutations

To generate the initial configuration of full-length PTEN mutants, we adopted the conformations of wild-type PTEN interacting with the membrane from previous studies (Jang *et al*., 2021). Explicit membrane simulations generated the fully relaxed wild-type proteins on an anionic lipid bilayer composed of DOPC:DOPS:PIP_2_:PIP_3_ (32:6:1:1 molar ratio). The wild-type sequence was modified to generate eight different PTEN mutants with each point mutation of Y68H, H93R, A126T, R130Q, G132D, R173C, F241S, and D252G. The anionic lipid bilayer with the same lipid compositions as in the wild-type system were reconstructed for the adopted mutant proteins. For all mutant systems, the initial configuration ensured that the PBD, phosphatase domain, and C2 domain were placed on the top of the bilayer surface without inserting the protein backbone into the bilayer, but the C-tail resided in bulky region without interacting with the lipid bilayer. Both PBD and C-tail were modeled as unstructured chains.

### Atomistic molecular dynamics simulations

MD simulations were performed on PTEN mutant systems using the updated CHARMM program with the modified all-atom force field (version 36m) (Brooks et al., 2009; Huang et al., 2017; Klauda et al., 2010). Our computational studies closely followed the same protocol as in our previous works (Grudzien et al., 2022; Haspel et al., 2021; Jang et al., 2016a; Jang et al., 2019; Jang et al., 2016b; Jang *et al*., 2021; Jang et al., 2020; Liao et al., 2020; Liu et al., 2022b; Liu et al., 2022c; Maloney et al., 2021; Maloney et al., 2022; Weako et al., 2021; Zhang et al., 2021a; Zhang et al., 2021b). Prior to productions runs, a series of minimization and dynamics cycles were performed for the solvents including ions and lipids with a harmonically restrained protein backbone until the solvent reached 310 K. Next, preequilibrium simulations with dynamic cycles were performed while gradually releasing the harmonic restraints on the backbones of PTEN mutants. The particle mesh Ewald (PME) method was used to calculate the long-range electrostatic interaction, and the van der Waals (vdW) interactions using switching functions with the twin range cutoff at 12 Å and 14 Å were calculated for the short-range interaction between atoms. In the production runs, the Nosé-Hoover Langevin piston control algorithm was used to sustain the pressure at 1 atm, and the Langevin thermostat method was employed to maintain the constant temperature at 310 K. The SHAKE algorithm was applied to constrain the motion of bonds involving hydrogen atoms. Simulations were performed for eight mutant systems each with 1 µs, and additional simulations for the same systems were also performed to check reproducibility. The production runs were performed with the NAMD parallel-computing code (Phillips et al., 2005) on a Biowulf cluster at the National Institutes of Health (Bethesda, MD). The result analysis was performed in the CHARMM program (Brooks *et al*., 2009). To determine the most populated conformation, the ensemble clustering in Chimera (Pettersen et al., 2004) was implemented to obtain the conformational representatives. The weighted implementation of suboptimal path (WISP) (Van Wart *et al*., 2014) algorithm was used to identify the allosteric signal propagation pathways through the protein. To observe conformational changes in proteins, the normal mode analysis (NMA) and principal component analysis (PCA) were conducted by the ProDy program (Bakan et al., 2011). In the analysis, averages were taken afterward discarding the first 200 ns trajectories.

### PTEN variants mapping and visualization

For variants mapping, eight PTEN mutations were considered: six in the phosphatase (Y68H, H93R, A126T, R130Q, G132D, and R173C) and two in the C2 (F241S and D252G) domains. The PTEN isoform structures were retrieved from the Release 42 (GRCh38.p13) of human genome on the GENCODE website (https://www.gencodegenes.org/human/). In total, we extracted isoform structures for seven PTEN isoforms. Only 5 isoforms, for which expression data was available, are shown in Figure 8. When we mapped PTEN variants to the isoforms, we only considered the exonic regions. The variants are grouped by the disease status (NDD vs. Non-NDD) and the two groups of variants are mapped and visualized separately. To perform the variants mapping, we used R language (v4.0.5) and RStudio. The Tidyverse package in R was used for data processing and data analysis. To generate the schematic figure for visualization of variants mapping results, we used the Gviz package in R.

### PTEN expression line plots

The expression profiles of PTEN isoforms were retrieved from the BrainSpan dataset which is an RNA-Seq datasets quantified at the gene and isoform levels and we downloaded it from PsychENCODE Knowledge Portal, PEC Capstone Collection, Synapse ID: syn8466658 (https://www.synapse.org/#!Synapse:syn12080241). The expression data was available for 5 out of 7 PTEN isoforms. For isoform expression level, transcripts per million (TPM) was used and log transformed. We used R language (v4.0.5) and RStudio to perform this analysis. The Tidyverse package in R was used for data processing, and the ggplot2 package in R was used for data visualization.

## Figure supplement legends

**Figure 1– figure supplement 1**. Lipid contact probability.

The probability of lipid contacts for PTEN residues for (A) the phosphatase mutations (Y68H, H93R, A126T, R130Q, G132D, and R173C) and (B) the C2 mutations (F241S and D252G). Also showing wild-type PTEN for comparison.

**Figure 1– figure supplement 2**. Helix tilt angle of PTEN.

Probability distribution functions of the helix tilt with respect to the bilayer normal for helices in the phosphatase domain of PTEN for (A) the phosphatase mutations (Y68H, H93R, A126T, R130Q, G132D, and R173C) and (B) the C2 mutations (F241S and D252G). Also showing wild-type PTEN for comparison.

**Figure 3–figure supplement 3**. Closed WPD loop conformation of PTEN H93R.

Violin plots representing the atomic pair distance between Cα of Asp92 in the WPD loop and Cα of Arg130 in the P loop for H93R and wild-type PTEN (left panel). The same plots for the distance between Cα of Arg93 (His93 for wild-type PTEN) in the WPD loop and Cα of Arg130 in the P loop for H93R (right panel).

**Figure 8–figure supplement 4**. The principal component analysis (PCA).

The projection of the first two principal components, PC1 and PC2, for the PTEN mutations, Y68H, H93R, A126T, R130Q, G132D, R173C, F241S, and D252G, and wild-type PTEN.

## Acknowledgements

LMI was supported by R01MH109885 and R01MH108528. This project has been funded in whole or in part with federal funds from the National Cancer Institute, National Institutes of Health, under contract HHSN261201500003I. The content of this publication does not necessarily reflect the views or policies of the Department of Health and Human Services, nor does mention of trade names, commercial products, or organizations imply endorsement by the U.S. Government. This Research was supported [in part] by the Intramural Research Program of the NIH, National Cancer Institute, Center for Cancer Research. All simulations had been performed using the high-performance computational facilities of the Biowulf PC/Linux cluster at the National Institutes of Health, Bethesda, MD (https://hpc.nih.gov/).

## Author contributions

H.J. built models and ran/analyzed molecular dynamics simulations. J.C. and L.M.I collected genomic data and retrieved BrainSpan dataset. H.J. wrote the initial draft, and L.M.I and R.N. edited the manuscript. R.N. supervised the project.

## References

Abkevich, V.I., Gutin, A.M., and Shakhnovich, E.I. 1995. Impact of local and non-local interactions on thermodynamics and kinetics of protein folding. J Mol Biol 252, 460–471. doi: 10.1006/jmbi.1995.0511.

Achterberg, J., Collerrain, I., and Craig, P. 1978. A possible relationship between cancer, mental retardation and mental disorders. Soc Sci Med (1967) 12, 135-139. doi.

Alvarez-Garcia, V., Tawil, Y., Wise, H.M., and Leslie, N.R. 2019. Mechanisms of PTEN loss in cancer: It’s all about diversity. Semin Cancer Biol 59, 66–79. doi: 10.1016/j.semcancer.2019.02.001.

Bakan, A., Meireles, L.M., and Bahar, I. 2011. ProDy: protein dynamics inferred from theory and experiments. Bioinformatics 27, 1575–1577. doi: 10.1093/bioinformatics/btr168.

Bolduc, D., Rahdar, M., Tu-Sekine, B., Sivakumaren, S.C., Raben, D., Amzel, L.M., Devreotes, P., Gabelli, S.B., and Cole, P. 2013. Phosphorylation-mediated PTEN conformational closure and deactivation revealed with protein semisynthesis. Elife 2, e00691. doi: 10.7554/eLife.00691.

Bonneau, D., and Longy, M. 2000. Mutations of the human PTEN gene. Hum Mutat 16, 109–122. doi: 10.1002/1098-1004(200008)16:2<109::AID-HUMU3>3.0.CO;2-0.

Brandao, T.A., Johnson, S.J., and Hengge, A.C. 2012. The molecular details of WPD-loop movement differ in the protein-tyrosine phosphatases YopH and PTP1B. Arch Biochem Biophys 525, 53–59. doi: 10.1016/j.abb.2012.06.002.

Brooks, B.R., Brooks, C.L., 3rd, Mackerell, A.D., Jr., Nilsson, L., Petrella, R.J., Roux, B., Won, Y., Archontis, G., Bartels, C., Boresch, S., Caflisch, A., Caves, L., Cui, Q., Dinner, A.R., Feig, M., Fischer, S., Gao, J., Hodoscek, M., Im, W., Kuczera, K., Lazaridis, T., Ma, J., Ovchinnikov, V., Paci, E., Pastor, R.W., Post, C.B., Pu, J.Z., Schaefer, M., Tidor, B., Venable, R.M., Woodcock, H.L., Wu, X., Yang, W., York, D.M., and Karplus, M. 2009. CHARMM: the biomolecular simulation program. J Comput Chem 30, 1545–1614. doi: 10.1002/jcc.21287.

Busch, R.M., Srivastava, S., Hogue, O., Frazier, T.W., Klaas, P., Hardan, A., Martinez-Agosto, J.A., Sahin, M., Eng, C., and Developmental Synaptopathies, C. 2019. Neurobehavioral phenotype of autism spectrum disorder associated with germline heterozygous mutations in PTEN. Transl Psychiatry 9, 253. doi: 10.1038/s41398-019-0588-1.

Butler, M.G., Dasouki, M.J., Zhou, X.P., Talebizadeh, Z., Brown, M., Takahashi, T.N., Miles, J.H., Wang, C.H., Stratton, R., Pilarski, R., and Eng, C. 2005. Subset of individuals with autism spectrum disorders and extreme macrocephaly associated with germline PTEN tumour suppressor gene mutations. J Med Genet 42, 318–321. doi: 10.1136/jmg.2004.024646.

Buxbaum, J.D., Cai, G., Chaste, P., Nygren, G., Goldsmith, J., Reichert, J., Anckarsater, H., Rastam, M., Smith, C.J., Silverman, J.M., Hollander, E., Leboyer, M., Gillberg, C., Verloes, A., and Betancur, C. 2007. Mutation screening of the PTEN gene in patients with autism spectrum disorders and macrocephaly. Am J Med Genet B Neuropsychiatr Genet 144B, 484–491. doi: 10.1002/ajmg.b.30493.

Chao, J.T., Hollman, R., Meyers, W.M., Meili, F., Matreyek, K.A., Dean, P., Fowler, D.M., Haas, K., Roskelley, C.D., and Loewen, C.J.R. 2020. A Premalignant Cell-Based Model for Functionalization and Classification of PTEN Variants. Cancer Res 80, 2775–2789. doi: 10.1158/0008-5472.CAN-19-3278.

Chau, K.K., Zhang, P., Urresti, J., Amar, M., Pramod, A.B., Chen, J., Thomas, A., Corominas, R., Lin, G.N., and Iakoucheva, L.M. 2021. Full-length isoform transcriptome of the developing human brain provides further insights into autism. Cell Rep 36, 109631. doi: 10.1016/j.celrep.2021.109631.

Cummings, K., Watkins, A., Jones, C., Dias, R., and Welham, A. 2022. Behavioural and psychological features of PTEN mutations: a systematic review of the literature and meta-analysis of the prevalence of autism spectrum disorder characteristics. J Neurodev Disord 14, 1. doi: 10.1186/s11689-021-09406-w.

Dempsey, D.R., Viennet, T., Iwase, R., Park, E., Henriquez, S., Chen, Z., Jeliazkov, J.R., Palanski, B.A., Phan, K.L., Coote, P., Gray, J.J., Eck, M.J., Gabelli, S.B., Arthanari, H., and Cole, P.A. 2021. The structural basis of PTEN regulation by multi-site phosphorylation. Nat Struct Mol Biol 28, 858–868. doi: 10.1038/s41594-021-00668-5.

Fricano-Kugler, C.J., Getz, S.A., Williams, M.R., Zurawel, A.A., DeSpenza, T., Jr., Frazel, P.W., Li, M., O’Malley, A.J., Moen, E.L., and Luikart, B.W. 2018. Nuclear Excluded Autism-Associated Phosphatase and Tensin Homolog Mutations Dysregulate Neuronal Growth. Biol Psychiatry 84, 265–277. doi: 10.1016/j.biopsych.2017.11.025.

Georgescu, M.M. 2010. PTEN Tumor Suppressor Network in PI3K-Akt Pathway Control. Genes Cancer 1, 1170–1177. doi: 10.1177/1947601911407325.

Grudzien, P., Jang, H., Leschinsky, N., Nussinov, R., and Gaponenko, V. 2022. Conformational Dynamics Allows Sampling of an “Active-like” State by Oncogenic K-Ras-GDP. J Mol Biol 434, 167695. doi: 10.1016/j.jmb.2022.167695.

Han, S.Y., Kato, H., Kato, S., Suzuki, T., Shibata, H., Ishii, S., Shiiba, K., Matsuno, S., Kanamaru, R., and Ishioka, C. 2000. Functional evaluation of PTEN missense mutations using in vitro phosphoinositide phosphatase assay. Cancer Res 60, 3147-3151. doi.

Haspel, N., Jang, H., and Nussinov, R. 2021. Active and Inactive Cdc42 Differ in Their Insert Region Conformational Dynamics. Biophys J 120, 306–318. doi: 10.1016/j.bpj.2020.12.007.

He, X., Ni, Y., Wang, Y., Romigh, T., and Eng, C. 2011. Naturally occurring germline and tumor-associated mutations within the ATP-binding motifs of PTEN lead to oxidative damage of DNA associated with decreased nuclear p53. Hum Mol Genet 20, 80–89. doi: 10.1093/hmg/ddq434.

Henager, S.H., Chu, N., Chen, Z., Bolduc, D., Dempsey, D.R., Hwang, Y., Wells, J., and Cole, P.A. 2016. Enzyme-catalyzed expressed protein ligation. Nat Methods 13, 925–927. doi: 10.1038/nmeth.4004.

Huang, J., Rauscher, S., Nawrocki, G., Ran, T., Feig, M., de Groot, B.L., Grubmuller, H., and MacKerell, A.D., Jr. 2017. CHARMM36m: an improved force field for folded and intrinsically disordered proteins. Nat Methods 14, 71–73. doi: 10.1038/nmeth.4067.

Jang, H., Banerjee, A., Chavan, T.S., Lu, S., Zhang, J., Gaponenko, V., and Nussinov, R. 2016a. The higher level of complexity of K-Ras4B activation at the membrane. FASEB J 30, 1643–1655. doi: 10.1096/fj.15-279091.

Jang, H., Banerjee, A., Marcus, K., Makowski, L., Mattos, C., Gaponenko, V., and Nussinov, R. 2019. The Structural Basis of the Farnesylated and Methylated KRas4B Interaction with Calmodulin. Structure 27, 1647–1659 e1644. doi: 10.1016/j.str.2019.08.009.

Jang, H., Muratcioglu, S., Gursoy, A., Keskin, O., and Nussinov, R. 2016b. Membrane-associated Ras dimers are isoform-specific: K-Ras dimers differ from H-Ras dimers. Biochem J 473, 1719–1732. doi: 10.1042/BCJ20160031.

Jang, H., Smith, I.N., Eng, C., and Nussinov, R. 2021. The mechanism of full activation of tumor suppressor PTEN at the phosphoinositide-enriched membrane. iScience 24, 102438. doi: 10.1016/j.isci.2021.102438.

Jang, H., Zhang, M., and Nussinov, R. 2020. The quaternary assembly of KRas4B with Raf-1 at the membrane. Comput Struct Biotechnol J 18, 737–748. doi: 10.1016/j.csbj.2020.03.018.

Kang, H.J., Kawasawa, Y.I., Cheng, F., Zhu, Y., Xu, X., Li, M., Sousa, A.M., Pletikos, M., Meyer, K.A., Sedmak, G., Guennel, T., Shin, Y., Johnson, M.B., Krsnik, Z., Mayer, S., Fertuzinhos, S., Umlauf, S., Lisgo, S.N., Vortmeyer, A., Weinberger, D.R., Mane, S., Hyde, T.M., Huttner, A., Reimers, M., Kleinman, J.E., and Sestan, N. 2011. Spatio-temporal transcriptome of the human brain. Nature 478, 483–489. doi: 10.1038/nature10523.

Klauda, J.B., Venable, R.M., Freites, J.A., O’Connor, J.W., Tobias, D.J., Mondragon-Ramirez, C., Vorobyov, I., MacKerell, A.D., Jr., and Pastor, R.W. 2010. Update of the CHARMM all-atom additive force field for lipids: validation on six lipid types. J Phys Chem B 114, 7830–7843. doi: 10.1021/jp101759q.

Koboldt, D.C., Miller, K.E., Miller, A.R., Bush, J.M., McGrath, S., Leraas, K., Crist, E., Fair, S., Schwind, W., Wijeratne, S., Fitch, J., Leonard, J., Shaikhouni, A., Hester, M.E., Magrini, V., Ho, M.L., Pierson, C.R., Wilson, R.K., Ostendorf, A.P., Mardis, E.R., and Bedrosian, T.A. 2021. PTEN somatic mutations contribute to spectrum of cerebral overgrowth. Brain 144, 2971–2978. doi: 10.1093/brain/awab173.

Kotelevets, L., Trifault, B., Chastre, E., and Scott, M.G.H. 2020. Posttranslational Regulation and Conformational Plasticity of PTEN. Cold Spring Harb Perspect Med 10. doi: 10.1101/cshperspect.a036095.

Lee, J.O., Yang, H., Georgescu, M.M., Di Cristofano, A., Maehama, T., Shi, Y., Dixon, J.E., Pandolfi, P., and Pavletich, N.P. 1999. Crystal structure of the PTEN tumor suppressor: implications for its phosphoinositide phosphatase activity and membrane association. Cell 99, 323–334. doi: 10.1016/s0092-8674(00)81663-3.

Li, M., Santpere, G., Imamura Kawasawa, Y., Evgrafov, O.V., Gulden, F.O., Pochareddy, S., Sunkin, S.M., Li, Z., Shin, Y., Zhu, Y., Sousa, A.M.M., Werling, D.M., Kitchen, R.R., Kang, H.J., Pletikos, M., Choi, J., Muchnik, S., Xu, X., Wang, D., Lorente-Galdos, B., Liu, S., Giusti-Rodriguez, P., Won, H., de Leeuw, C.A., Pardinas, A.F., BrainSpan, C., Psych, E.C., Psych, E.D.S., Hu, M., Jin, F., Li, Y., Owen, M.J., O’Donovan, M.C., Walters, J.T.R., Posthuma, D., Reimers, M.A., Levitt, P., Weinberger, D.R., Hyde, T.M., Kleinman, J.E., Geschwind, D.H., Hawrylycz, M.J., State, M.W., Sanders, S.J., Sullivan, P.F., Gerstein, M.B., Lein, E.S., Knowles, J.A., and Sestan, N. 2018. Integrative functional genomic analysis of human brain development and neuropsychiatric risks. Science 362. doi: 10.1126/science.aat7615.

Liao, T.J., Jang, H., Fushman, D., and Nussinov, R. 2020. SOS1 interacts with Grb2 through regions that induce closed nSH3 conformations. J Chem Phys 153, 045106. doi: 10.1063/5.0013926.

Liu, Q., Adami, H.O., Reichenberg, A., Kolevzon, A., Fang, F., and Sandin, S. 2021. Cancer risk in individuals with intellectual disability in Sweden: A population-based cohort study. PLoS Med 18, e1003840. doi: 10.1371/journal.pmed.1003840.

Liu, Q., Yin, W., Meijsen, J.J., Reichenberg, A., Gadin, J.R., Schork, A.J., Adami, H.O., Kolevzon, A., Sandin, S., and Fang, F. 2022a. Cancer risk in individuals with autism spectrum disorder. Ann Oncol 33, 713–719. doi: 10.1016/j.annonc.2022.04.006.

Liu, Y., Jang, H., Zhang, M., Tsai, C.J., Maloney, R., and Nussinov, R. 2022b. The structural basis of BCR-ABL recruitment of GRB2 in chronic myelogenous leukemia. Biophys J 121, 2251–2265. doi: 10.1016/j.bpj.2022.05.030.

Liu, Y., Zhang, M., Tsai, C.J., Jang, H., and Nussinov, R. 2022c. Allosteric regulation of autoinhibition and activation of c-Abl. Comput Struct Biotechnol J 20, 4257–4270. doi: 10.1016/j.csbj.2022.08.014.

Malaney, P., Pathak, R.R., Xue, B., Uversky, V.N., and Dave, V. 2013. Intrinsic disorder in PTEN and its interactome confers structural plasticity and functional versatility. Sci Rep 3, 2035. doi: 10.1038/srep02035.

Maloney, R.C., Zhang, M., Jang, H., and Nussinov, R. 2021. The mechanism of activation of monomeric B-Raf V600E. Comput Struct Biotechnol J 19, 3349–3363. doi: 10.1016/j.csbj.2021.06.007.

Maloney, R.C., Zhang, M., Liu, Y., Jang, H., and Nussinov, R. 2022. The mechanism of activation of MEK1 by B-Raf and KSR1. Cell Mol Life Sci 79, 281. doi: 10.1007/s00018-022-04296-0.

Marsh, D.J., Theodosopoulos, G., Howell, V., Richardson, A.L., Benn, D.E., Proos, A.L., Eng, C., and Robinson, B.G. 2001. Rapid mutation scanning of genes associated with familial cancer syndromes using denaturing high-performance liquid chromatography. Neoplasia 3, 236–244. doi: 10.1038/sj.neo.7900154.

Meng, Z., Jia, L.F., and Gan, Y.H. 2016. PTEN activation through K163 acetylation by inhibiting HDAC6 contributes to tumour inhibition. Oncogene 35, 2333–2344. doi: 10.1038/onc.2015.293.

Mingo, J., Rodriguez-Escudero, I., Luna, S., Fernandez-Acero, T., Amo, L., Jonasson, A.R., Zori, R.T., Lopez, J.I., Molina, M., Cid, V.J., and Pulido, R. 2018. A pathogenic role for germline PTEN variants which accumulate into the nucleus. Eur J Hum Genet 26, 1180–1187. doi: 10.1038/s41431-018-0155-x.

Morris-Rosendahl, D.J., and Crocq, M.A. 2020. Neurodevelopmental disorders-the history and future of a diagnostic concept. Dialogues Clin Neurosci 22, 65–72. doi: 10.31887/DCNS.2020.22.1/macrocq.

Nanda, H., Heinrich, F., and Losche, M. 2015. Membrane association of the PTEN tumor suppressor: neutron scattering and MD simulations reveal the structure of protein-membrane complexes. Methods 77-78, 136–146. doi: 10.1016/j.ymeth.2014.10.014.

Nordentoft, M., Plana-Ripoll, O., and Laursen, T.M. 2021. Cancer and schizophrenia. Curr Opin Psychiatry 34, 260–265. doi: 10.1097/YCO.0000000000000697.

Nussinov, R., Tsai, C.J., and Jang, H. 2022a. Allostery, and how to define and measure signal transduction. Biophys Chem 283, 106766. doi: 10.1016/j.bpc.2022.106766.

Nussinov, R., Tsai, C.J., and Jang, H. 2022b. How can same-gene mutations promote both cancer and developmental disorders? Sci Adv 8, eabm2059. doi: 10.1126/sciadv.abm2059.

Nussinov, R., Tsai, C.J., and Jang, H. 2022c. Neurodevelopmental disorders, immunity, and cancer are connected. iScience 25, 104492. doi: 10.1016/j.isci.2022.104492.

Nussinov, R., Tsai, C.J., and Jang, H. 2022d. A New View of Activating Mutations in Cancer. Cancer Res 82, 4114–4123. doi: 10.1158/0008-5472.CAN-22-2125.

Pettersen, E.F., Goddard, T.D., Huang, C.C., Couch, G.S., Greenblatt, D.M., Meng, E.C., and Ferrin, T.E. 2004. UCSF Chimera--a visualization system for exploratory research and analysis. J Comput Chem 25, 1605–1612. doi: 10.1002/jcc.20084.

Phillips, J.C., Braun, R., Wang, W., Gumbart, J., Tajkhorshid, E., Villa, E., Chipot, C., Skeel, R.D., Kale, L., and Schulten, K. 2005. Scalable molecular dynamics with NAMD. J Comput Chem 26, 1781–1802. doi: 10.1002/jcc.20289.

Pilarski, R., Burt, R., Kohlman, W., Pho, L., Shannon, K.M., and Swisher, E. 2013. Cowden syndrome and the PTEN hamartoma tumor syndrome: systematic review and revised diagnostic criteria. J Natl Cancer Inst 105, 1607–1616. doi: 10.1093/jnci/djt277.

Post, K.L., Belmadani, M., Ganguly, P., Meili, F., Dingwall, R., McDiarmid, T.A., Meyers, W.M., Herrington, C., Young, B.P., Callaghan, D.B., Rogic, S., Edwards, M., Niciforovic, A., Cau, A., Rankin, C.H., O’Connor, T.P., Bamji, S.X., Loewen, C.J.R., Allan, D.W., Pavlidis, P., and Haas, K. 2020. Multi-model functionalization of disease-associated PTEN missense mutations identifies multiple molecular mechanisms underlying protein dysfunction. Nat Commun 11, 2073. doi: 10.1038/s41467-020-15943-0.

Rahdar, M., Inoue, T., Meyer, T., Zhang, J., Vazquez, F., and Devreotes, P.N. 2009. A phosphorylation-dependent intramolecular interaction regulates the membrane association and activity of the tumor suppressor PTEN. Proc Natl Acad Sci U S A 106, 480–485. doi: 10.1073/pnas.0811212106.

Redfern, R.E., Daou, M.C., Li, L., Munson, M., Gericke, A., and Ross, A.H. 2010. A mutant form of PTEN linked to autism. Protein Sci 19, 1948–1956. doi: 10.1002/pro.483.

Rodriguez-Escudero, I., Oliver, M.D., Andres-Pons, A., Molina, M., Cid, V.J., and Pulido, R. 2011. A comprehensive functional analysis of PTEN mutations: implications in tumor- and autism-related syndromes. Hum Mol Genet 20, 4132–4142. doi: 10.1093/hmg/ddr337.

Ross, A.H., and Gericke, A. 2009. Phosphorylation keeps PTEN phosphatase closed for business. Proc Natl Acad Sci U S A 106, 1297–1298. doi: 10.1073/pnas.0812473106.

Sansal, I., and Sellers, W.R. 2004. The biology and clinical relevance of the PTEN tumor suppressor pathway. J Clin Oncol 22, 2954–2963. doi: 10.1200/JCO.2004.02.141.

Serebriiskii, I.G., Pavlov, V., Tricarico, R., Andrianov, G., Nicolas, E., Parker, M.I., Newberg, J., Frampton, G., Meyer, J.E., and Golemis, E.A. 2022. Comprehensive characterization of PTEN mutational profile in a series of 34,129 colorectal cancers. Nat Commun 13, 1618. doi: 10.1038/s41467-022-29227-2.

Shan, L., Yu, J., He, Z., Chen, S., Liu, M., Ding, H., Xu, L., Zhao, J., Yang, A., and Jiang, H. 2020. Defining relative mutational difficulty to understand cancer formation. Cell Discov 6, 48. doi: 10.1038/s41421-020-0177-8.

Shenoy, S.S., Nanda, H., and Losche, M. 2012. Membrane association of the PTEN tumor suppressor: electrostatic interaction with phosphatidylserine-containing bilayers and regulatory role of the C-terminal tail. J Struct Biol 180, 394–408. doi: 10.1016/j.jsb.2012.10.003.

Singh, G., and Chan, A.M. 2011. Post-translational modifications of PTEN and their potential therapeutic implications. Curr Cancer Drug Targets 11, 536–547. doi: 10.2174/156800911795655930.

Song, M.S., Salmena, L., and Pandolfi, P.P. 2012. The functions and regulation of the PTEN tumour suppressor. Nat Rev Mol Cell Biol 13, 283–296. doi: 10.1038/nrm3330.

Spinelli, L., Black, F.M., Berg, J.N., Eickholt, B.J., and Leslie, N.R. 2015. Functionally distinct groups of inherited PTEN mutations in autism and tumour syndromes. J Med Genet 52, 128–134. doi: 10.1136/jmedgenet-2014-102803.

Strokach, A., and Kim, P.M. 2022. Deep generative modeling for protein design. Curr Opin Struct Biol 72, 226–236. doi: 10.1016/j.sbi.2021.11.008.

Tan, M.H., Mester, J., Peterson, C., Yang, Y., Chen, J.L., Rybicki, L.A., Milas, K., Pederson, H., Remzi, B., Orloff, M.S., and Eng, C. 2011. A clinical scoring system for selection of patients for PTEN mutation testing is proposed on the basis of a prospective study of 3042 probands. Am J Hum Genet 88, 42–56. doi: 10.1016/j.ajhg.2010.11.013.

Tan, M.H., Mester, J.L., Ngeow, J., Rybicki, L.A., Orloff, M.S., and Eng, C. 2012. Lifetime cancer risks in individuals with germline PTEN mutations. Clin Cancer Res 18, 400–407. doi: 10.1158/1078-0432.CCR-11-2283.

Tsou, H.C., Ping, X.L., Xie, X.X., Gruener, A.C., Zhang, H., Nini, R., Swisshelm, K., Sybert, V., Diamond, T.M., Sutphen, R., and Peacocke, M. 1998. The genetic basis of Cowden’s syndrome: three novel mutations in PTEN/MMAC1/TEP1. Hum Genet 102, 467–473. doi: 10.1007/s004390050723.

Tu, T., Chen, J., Chen, L., and Stiles, B.L. 2020. Dual-Specific Protein and Lipid Phosphatase PTEN and Its Biological Functions. Cold Spring Harb Perspect Med 10. doi: 10.1101/cshperspect.a036301.

Valtcheva, N., Lang, F.M., Noske, A., Samartzis, E.P., Schmidt, A.M., Bellini, E., Fink, D., Moch, H., Rechsteiner, M., Dedes, K.J., and Wild, P.J. 2017. Tracking the origin of simultaneous endometrial and ovarian cancer by next-generation sequencing - a case report. BMC Cancer 17, 66. doi: 10.1186/s12885-017-3054-6.

Van Wart, A.T., Durrant, J., Votapka, L., and Amaro, R.E. 2014. Weighted Implementation of Suboptimal Paths (WISP): An Optimized Algorithm and Tool for Dynamical Network Analysis. J Chem Theory Comput 10, 511–517. doi: 10.1021/ct4008603.

Waite, K.A., and Eng, C. 2002. Protean PTEN: form and function. Am J Hum Genet 70, 829–844. doi: 10.1086/340026.

Weako, J., Jang, H., Keskin, O., Nussinov, R., and Gursoy, A. 2021. The structural basis of Akt PH domain interaction with calmodulin. Biophys J 120, 1994–2008. doi: 10.1016/j.bpj.2021.03.018.

Xia, Q., Ali, S., Liu, L., Li, Y., Liu, X., Zhang, L., and Dong, L. 2020. Role of Ubiquitination in PTEN Cellular Homeostasis and Its Implications in GB Drug Resistance. Front Oncol 10, 1569. doi: 10.3389/fonc.2020.01569.

Xu, W., Yang, Z., Zhou, S.F., and Lu, N. 2014. Posttranslational regulation of phosphatase and tensin homolog (PTEN) and its functional impact on cancer behaviors. Drug Des Devel Ther 8, 1745–1751. doi: 10.2147/DDDT.S71061.

Yehia, L., Ni, Y., Sadler, T., Frazier, T.W., and Eng, C. 2022. Distinct metabolic profiles associated with autism spectrum disorder versus cancer in individuals with germline PTEN mutations. NPJ Genom Med 7, 16. doi: 10.1038/s41525-022-00289-x.

Yin, Y., and Shen, W.H. 2008. PTEN: a new guardian of the genome. Oncogene 27, 5443–5453. doi: 10.1038/onc.2008.241.

Zhang, M., Jang, H., Li, Z., Sacks, D.B., and Nussinov, R. 2021a. B-Raf autoinhibition in the presence and absence of 14-3-3. Structure 29, 768–777 e762. doi: 10.1016/j.str.2021.02.005.

Zhang, M., Maloney, R., Jang, H., and Nussinov, R. 2021b. The mechanism of Raf activation through dimerization. Chem Sci 12, 15609–15619. doi: 10.1039/d1sc03444h.

Zhang, Y., Park, J., Han, S.J., Yang, S.Y., Yoon, H.J., Park, I., Woo, H.A., and Lee, S.R. 2020. Redox regulation of tumor suppressor PTEN in cell signaling. Redox Biol 34, 101553. doi: 10.1016/j.redox.2020.101553.

