## Supplemental Figures for "How PTEN mutations degrade function at the membrane and life expectancy of carriers of mutations in the human brain"

**A** Phosphatase mutations

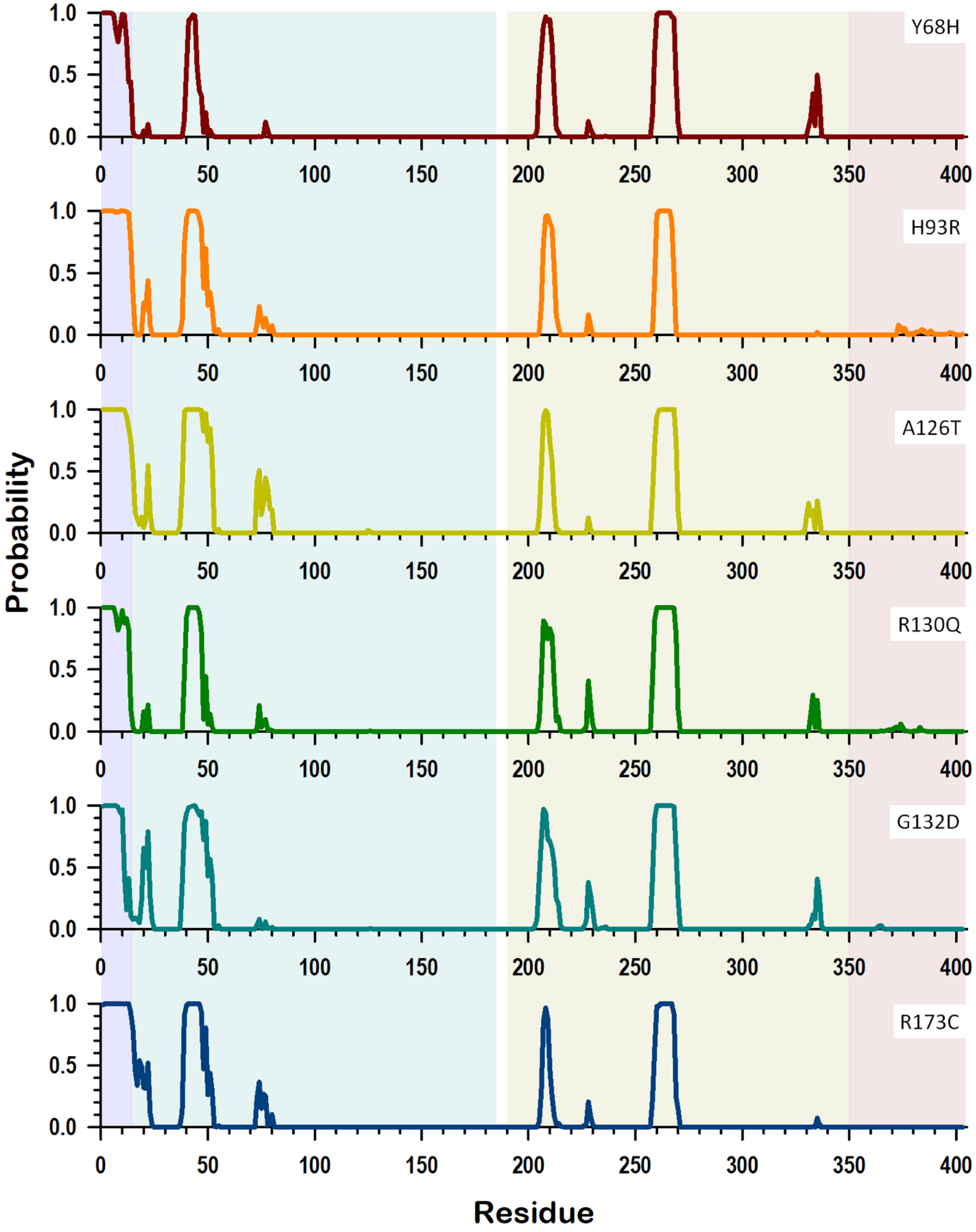

**B** C2 mutations

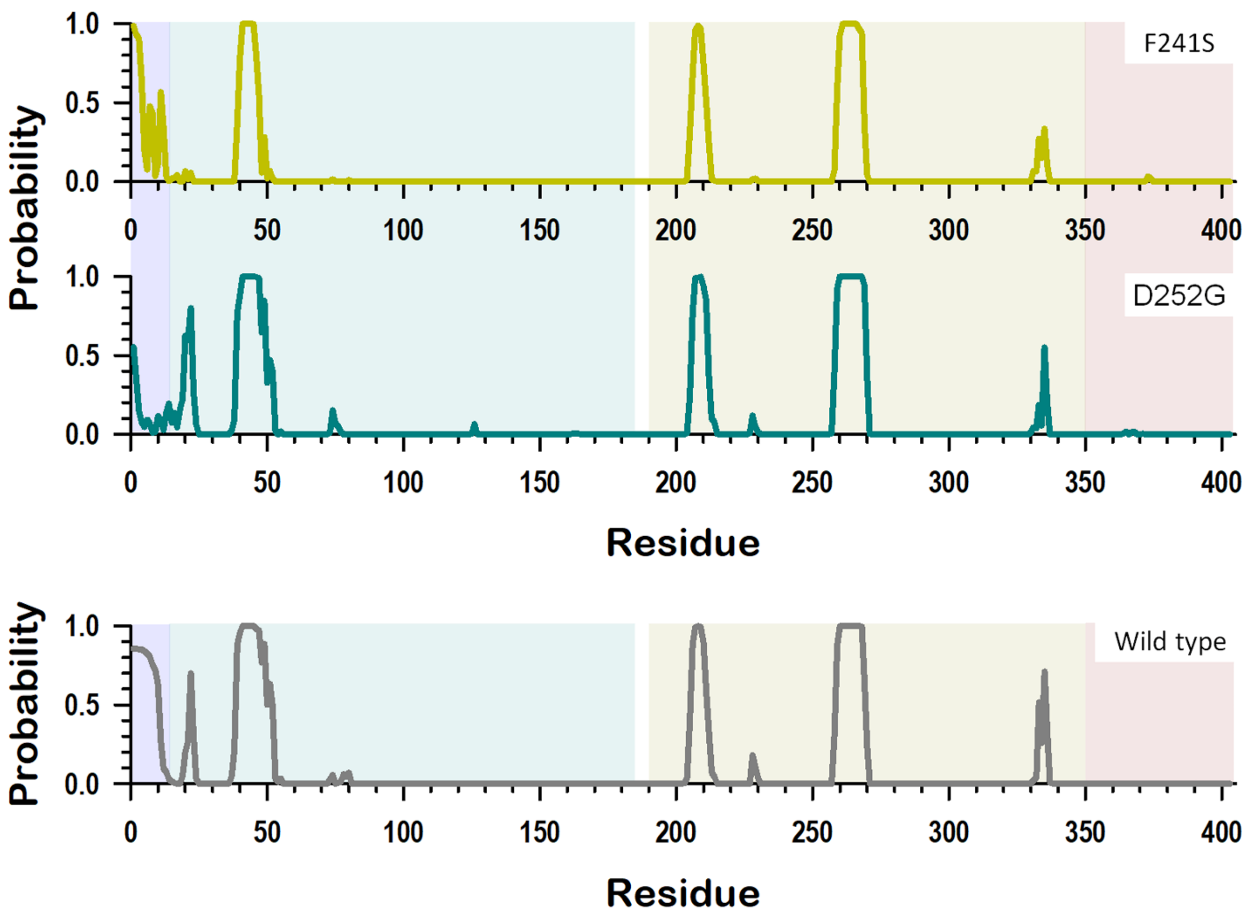


**Supplementary Figure 1.** Lipid contact probability. The probability of lipid contacts for PTEN residues for (A) the phosphatase mutations (Y68H, H93R, A126T, R130Q, G132D, and R173C) and (B) the C2 mutations (F241S and D252G). Also showing wild-type PTEN for comparison.

**A** Phosphatase mutations

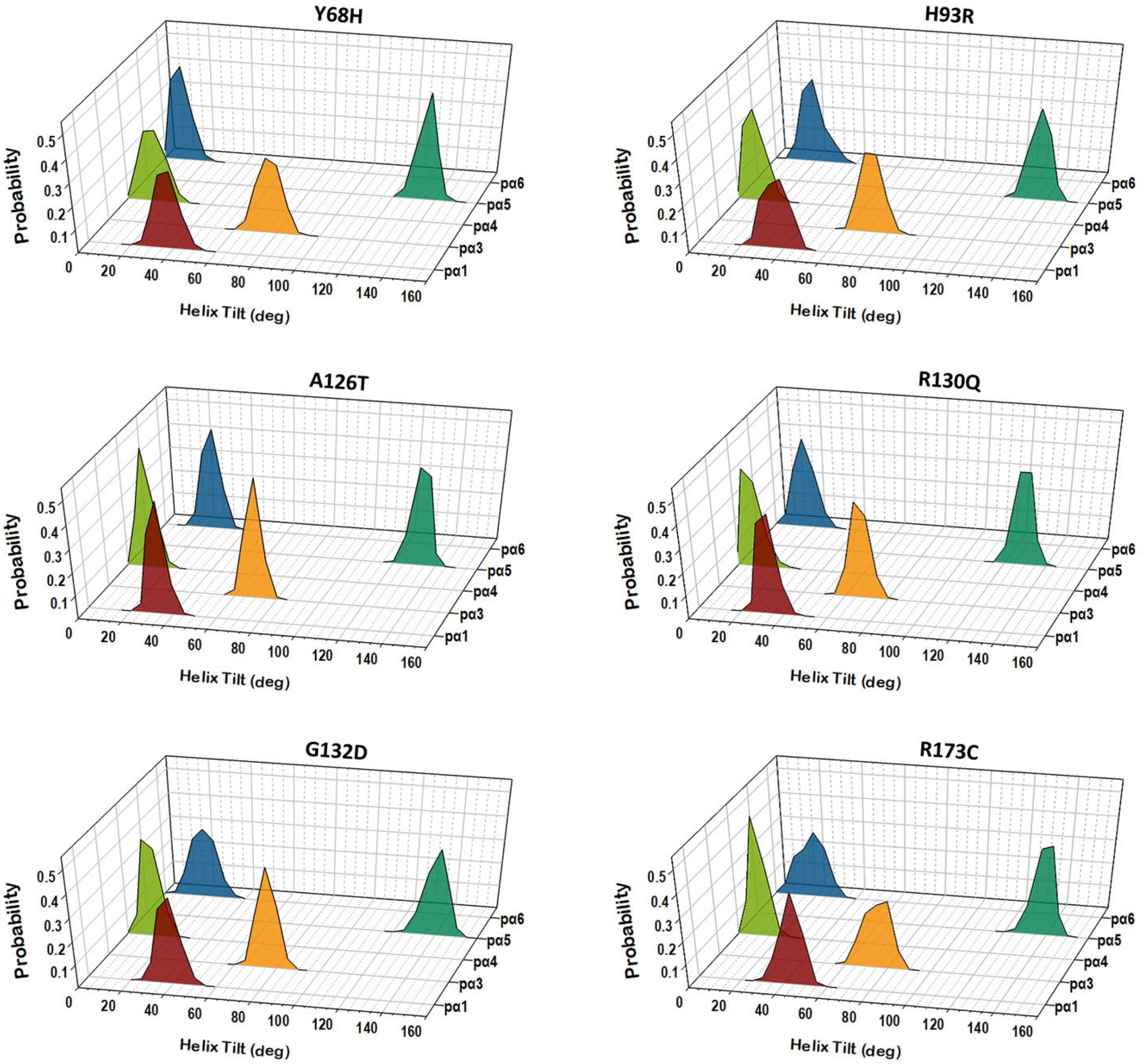

**B** C2 mutations

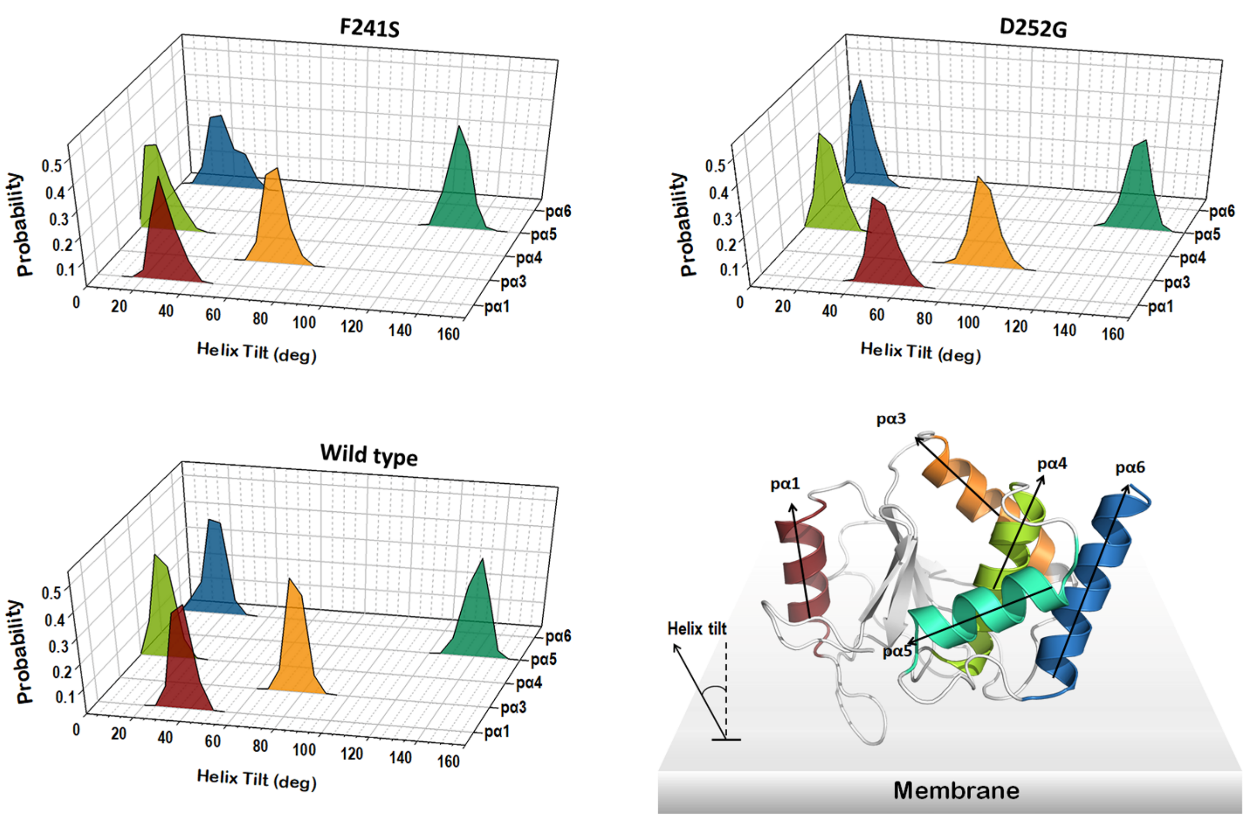


**Supplementary Figure 2.** Helix tilt angle of PTEN. Probability distribution functions of the helix tilt with respect to the bilayer normal for helices in the phosphatase domain of PTEN for (A) the phosphatase mutations (Y68H, H93R, A126T, R130Q, G132D, and R173C) and (B) the C2 mutations (F241S and D252G). Also showing wild-type PTEN for comparison.

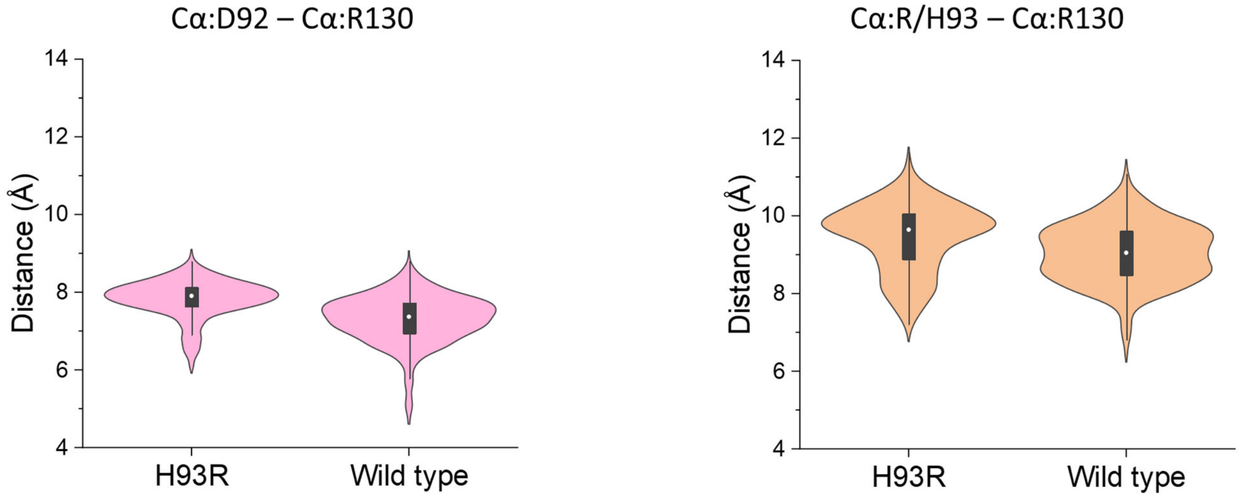


**Supplementary Figure 3.** Closed WPD loop conformation of PTEN H93R. Violin plots representing the atomic pair distance between Cα of Asp92 in the WPD loop and Cα of Arg130 in the P loop for H93R and wild-type PTEN (left panel). The same plots for the distance between Cα of Arg93 (His93 for wild-type PTEN) in the WPD loop and Cα of Arg130 in the P loop for H93R (right panel).

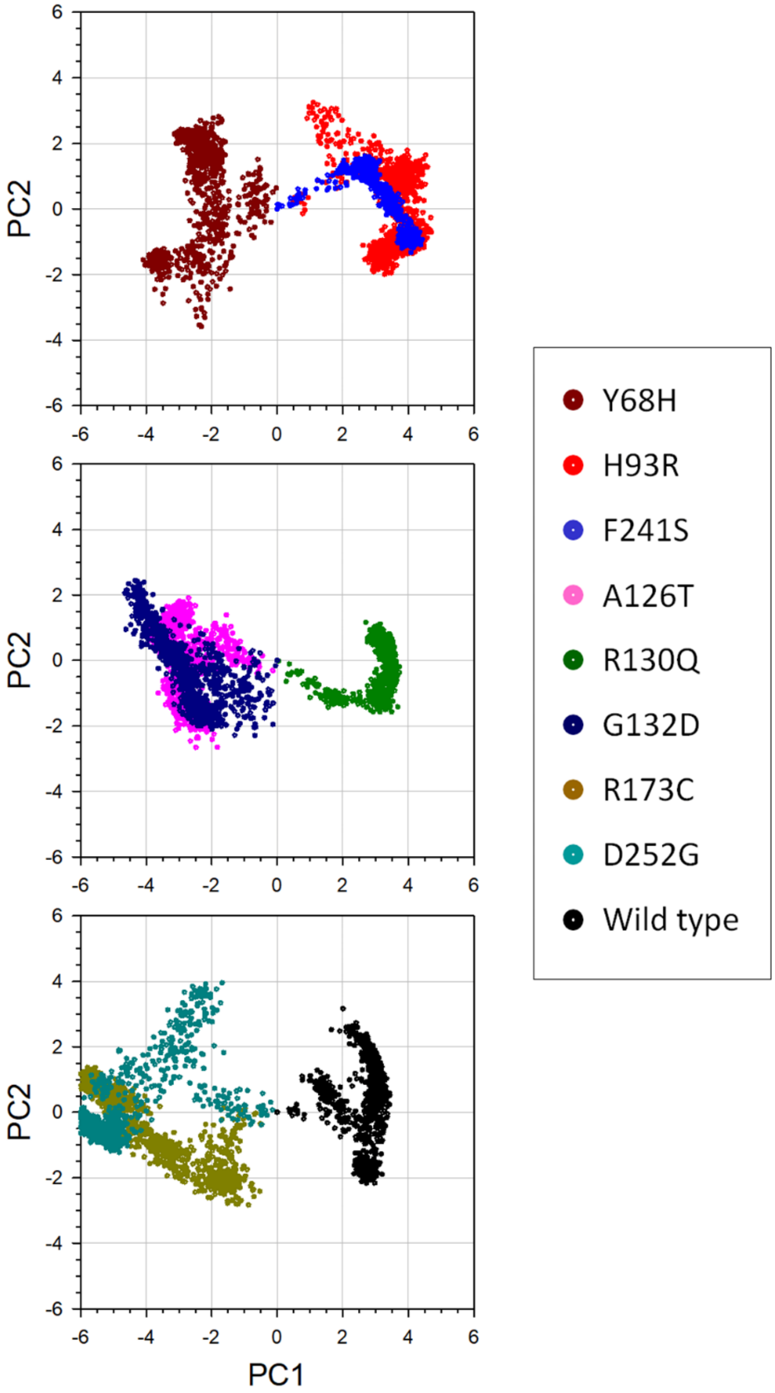


**Supplementary Figure 4.** The principal component analysis (PCA). The projection of the first two principal components, PC1 and PC2, for the PTEN mutations, Y68H, H93R, A126T, R130Q, G132D, R173C, F241S, and D252G, and wild-type PTEN.
